# TREM2 promotes glioma progression and angiogenesis mediated by microglia/brain macrophages

**DOI:** 10.1101/2023.05.18.540621

**Authors:** Xuezhen Chen, Yue Zhao, Yimin Huang, Kaichuan Zhu, Fan Zeng, Junyi Zhao, Huaqiu Zhang, Xinzhou Zhu, Helmut Kettenmann, Xianyuan Xiang

## Abstract

TREM2, a myeloid cell-specific signaling molecule, controls essential functions of microglia and impacts on the pathogenesis of Alzheimer’s disease and other neurodegenerative disorders. TREM2 is also highly expressed in tumor-associated macrophages and plays detrimental roles in an experimental mouse sarcoma model. Here we studied whether TREM2 influences glioma progression. We found a sex-dependent effect of TREM2: the glioma volume is significantly attenuated in TREM2- deficient male but not female mice injected with GL261-EGFP glioma cells. The accumulation of glioma-associated microglia/macrophages (GAMs) and vascularization is reduced in male TREM2-deficient mice. A transcriptomic analysis of glioma tissue revealed that TREM2 deficiency suppresses angiogenic genes and MHC clusters. In an organotypic slice model devoid of functional vascularization, the tumor size was not affected by TREM2-deficiency. In human resection samples from glioblastoma, TREM2 is upregulated in GAMs. Based on the TCGA and CGGA databases, the TREM2 expression levels are negatively correlated with survival. Thus, the TREM2-dependent crosstalk between GAMs and the vasculature formation promotes glioma growth.

**Graphic abstract:** TREM2-dependent crosstalk between glioma-associated microglia/macrophages and the vasculature formation promotes glioma growth in male glioma mouse model. Created with BioRender.com

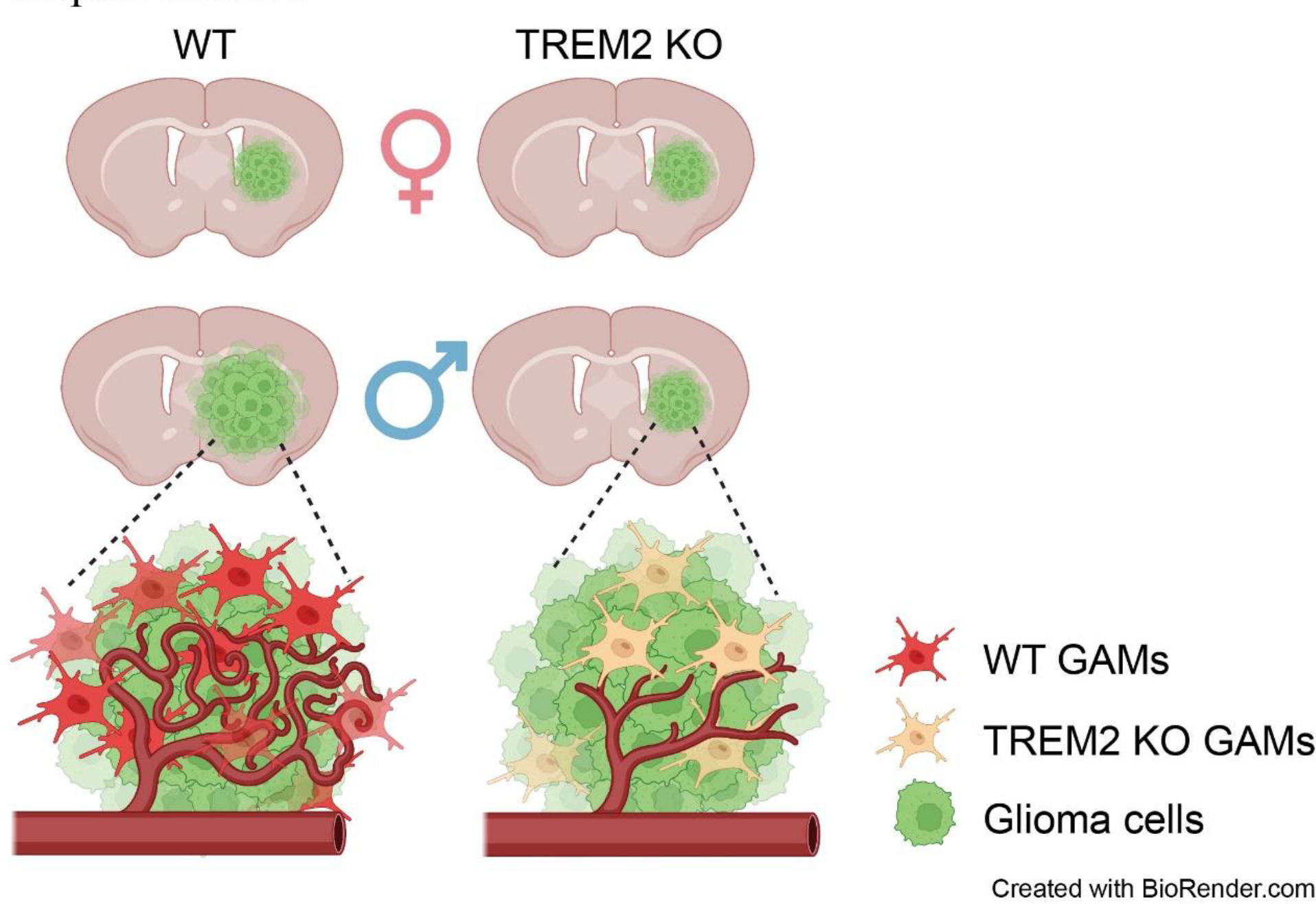

## Introduction

Glioma is the most common form of tumor within the central nervous system (CNS), and is classified into low-grade (II) and high-grade (III or IV) glioblastomas (GBM) (Gutmann and Kettenmann, 2019). GBM is the most lethal malignant brain tumor in adults; currently, no effective treatment is available. The incidence of GBM is higher in men than women, with a ratio of 1.6:1 (Davis, 2016; Ostrom et al., 2018b). With the standard treatments, namely surgical resection followed by radio-/chemotherapy, the median survival rates for GBM patients remains disappointedly low with a 5-year survival rate of 9.8% (Schaff and Mellinghoff, 2023), although the standard therapy is more effective in female than in male patients (Yang et al., 2019). Immunotherapy with checkpoint inhibitors has shown great success in various solid tumors, whereas less than 10% of GBM patients show long-term responses (Jackson et al., 2019). The complexity and heterogeneity of the brain tumor microenvironment (TME) is one of the main obstacles for developing novel therapeutic strategies (Dumas et al., 2020). In addition to cancer cells, TME contains various non-cancerous cell types, including immune and endothelial cells (Wu et al., 2021). The communication and reciprocal connections among these cells create an immuno-suppressive environment and actively contribute to GBM progression (Charles et al., 2012; Hambardzumyan et al., 2016; Gutmann and Kettenmann, 2019; Liu et al., 2022).

The majority of immune cells in the GBM environment is represented by the myeloid cell population, composed of brain resident microglia and infiltrated macrophages, termed glioma-associated microglia/macrophages (GAMs). They often comprise up to 30%-50% of the tumor mass (Gutmann and Kettenmann, 2019) and outnumber infiltrating T cells (Lim et al., 2018), providing an immunosuppressive and pro-tumorigenic environment. The glioma environment convert GAMs into tumor-promoting cells, which enhances glioma cell proliferation, degrades the extracellular matrix, promotes angiogenesis, and induces T-cell apoptosis (Roesch et al., 2018; Gutmann and Kettenmann, 2019). Therefore, GAMs have become attractive novel targets for glioma therapy. However, GAMs depletion using colony-stimulating factor- 1 receptor (CSF-1R) inhibitors failed to improve the clinical outcome (Binnewies et al., 2021). Consequently, more precise GAMs-regulating targets should be identified to better understand the complex interaction of glioma and glioma associated cells including the vascular system.

Triggering receptor expressed on myeloid cells 2 (TREM2) is exclusively expressed in the myeloid cell lineage, regulating important myeloid cell functions, including proliferation (Schlepckow et al., 2020), migration (Mazaheri et al., 2017), and phagocytosis (Hsieh et al., 2009; Kleinberger et al., 2014; Xiang et al., 2016). TREM2 restrains cytokine production in macrophages in response to LPS stimulation (Turnbull et al., 2006). TREM2 is upregulated in macrophages associated with human and mouse tumors. In a sarcoma mouse model, inhibition of TREM2 signaling restricts tumor growth and sensitizes the response to anti-PD1 immunotherapy (Katzenelenbogen et al., 2020; Molgora et al., 2020). A humanized antibody targeting TREM2 is currently tested in phase I clinical trials for solid tumors (Binnewies et al., 2021). While there is good evidence for the involvement of TREM2 in other solid tumors, its role in glioma remains to be determined.

In this study, we used the orthotopic GL261 mouse glioma model (Haddad et al., 2021) *in vivo* and the well-established organotypic brain slice (OBS) glioma model (Hu et al., 2015; Huang et al., 2020) *ex vivo* to explore the functional role of TREM2 in the context of GBM progression. We found that the crosstalk between microglia and the vasculature mediated by TREM2 has an impact on glioma progression. As a result, the TREM2- dependent vascularization is correlated with larger glioma volume.

## Methods and materials

### Animals

The TREM2 KO (Turnbull et al., 2006) line was kindly provided by Dr. Marco Colonna at the Washington University School of Medicine, St. Louis. The line is maintained at heterozygous breeding and WT or TREM2 KO littermates were used from the offspring. All experiments were approved by the Shenzhen Institutes of Advanced Technology, Chinese Academy of Sciences Research Committee, and all experimental procedures involving animals were carried out in strict accordance with the Research Committee’s animal user guidelines. Food and water were provided ad libitum. Male and female mice (10-12 weeks) were used in this study.

### Cell culture

The murine glioma cell line GL261 (American Type Culture Collection) was cultured in DMEM supplemented with 10% FCS (Lonsera, Suzhou, China), 50 units/ml penicillin-streptomycin, and 200 mM glutamine (Invitrogen, Carlsbad, USA). GL261 were transduced with lentivirus expressing enhanced green fluorescent protein (EGFP). For primary microglia cultures, P1-P3 mice were used. After extraction of the brains, meninges and cerebellum were removed and brains were put on ice with Hanks balanced salt solution (HBSS, Beyotime Biotechnology, Shanghai, China), subsequently minced into small pieces before adding trypsin (10 mg/mL, Beyotime Biotechnology) and DNase (0.5 mg/mL, Beyotime Biotechnology) in PBS. After 2–3 min of incubation, the reaction was blocked by adding Dulbecco’s Modified Eagle Medium (DMEM) containing 10% fetal bovine serum (FBS). After removal of the medium, 10 U of DNase was added and the cells were mechanically dissociated. The dissociated cell suspension was centrifuged for 10 min at 300g at 4°C. The pellet was resuspended in DMEM supplements (10% heat-inactivated FCS, Lonsera), 50 units/ml penicillin-streptomycin, and 200 mM glutamine (Invitrogen) and plated (cells from 2 brains/flask) in poly-L-lysine-coated T75 flasks (Corning, New York, USA). A complete change of medium was performed 24 hours after plating to remove excess debris. Thereafter, half of the medium was changed every two days. After 9-10 days in culture, primary microglia were collected from supernatant and plated at a density of 3× 10^5^ in one well of a 12-wall plate. The supernatant of the cultivated primary microglia was collected 3 days after plating. The microglia conditioned medium was stored at -20^0^C until further usage.

### *In vivo* glioma model

Ten- to twelve-week-old WT and TREM2 KO mice were used for the *in vivo* studies to investigate glioma progression. Mice were anesthetized, immobilized, and mounted onto a stereotactic frame in the flat-skull position (Digital Stereotaxic Instruments, 68018 RWD Life Science, Shenzhen, China). After skin incision, the skull was carefully drilled at the point located at 1 mm anterior and 1.8 mm lateral to the bregma with Hamilton tip. A 2 μl syringe with a blunt tip (Mikroliterspritze 7001 N, Hamilton, Reno, USA) was inserted to a depth of 4.5 mm and retracted to a depth of 4 mm from the skull into the right caudate putamen. Then 0.5 μl of a cell suspension containing 1× 10^4^ GL261-EGFP glioma cells were slowly injected into the brain within 2.5 min. The needle was then slowly retracted from the injection canal, and the skin was sutured with a surgical sewing cone (RWD Life Science). After surgery, the mice were kept on a 37 ^0^C heated pat until awake. Mice were monitored daily and perfused with paraformaldehyde (PFA) 21 days after injection, and brain samples were collected for the immunohistochemistry and RNA sequencing.

### Tumor Size Measurements

After animal perfusion, brains were fixed with 4% PFA in PBS (Beyotime Biotechnology) overnight at 4 ℃ and dehydrated with 30% sucrose for 48 h at 4 ℃. To prepare brain sections, we cut brains into 40 μm sections, collected every 12^th^ slice and mounted it on a glass slide. Other slices were put in cryoprotectant solution for immunofluorescence staining and inspection. The EGFP fluorescence signal of the GL261 glioma cells was excited at a wavelength 488 nm and visualized at of 507 nm analyzed by a Zeiss LSM apotome microscope (Carl Zeiss, Oberkochen, Germany) with a 10×objective. The glioma area in individual slices was determined based on the fluorescence area analyzed with ImageJ software. Tumor volumes were determined by H&E staining every 12th slice according to the Cavalieri principle (Hu et al., 2015; Huang et al., 2020).

### Mice survival assay

For survival analysis, an independent cohort of mice was used. The GL261-EGFP- injected mice were monitored daily from the 21 days until the terminal endpoint. The disease progression was scored from 0 to 20 based on the severity of following criteria: loss of body weight, hunchback, low reaction to stimulation, tarnished fur, or movement problem (normal = 0, slight burden = 5, medium burden = 10, severe burden = 20). When the sum of scores of these criteria is equal or larger than 20, the animals were sacrificed.

### Organotypic brain slice (OBS) cultures

OBS were prepared as described previously (Huang et al., 2020) with minor modifications. P5-8 WT and TREM2 KO mice were sacrificed, brains were taken out, and placed in cold HBSS (Beyotime Biotechnology). The left and right hemisphere was separated and cut coronally into 250 μm thick slices using McIlwain tissue chopper (Ted Pella Inc, Redding, USA) under a laminar hood. Six to eight slices were transferred individually into a Millicell Cell Culture Insert, 30 mm, hydrophilic polytetrafluoroethylene polymer, 0.4 µm (PICM0RG50 Millipore, Darmstadt, Germany) in 6-well plate with culture medium containing 50% DMEM, 25% Basal Medium Eagle (BME), 25% heat inactivated horse serum, 10mM HEPES, 10mM Tris, 0.1% Glucose, 2nM Glutamax, NaOH PH7.4 and Insulin 5 μg/ml (Invitrogen). Culture medium was changed every two days. After 5 days of culture maintenance, 5000 GL261-EGFP cells in a volume of 0.1μl were inoculated into the brain slices using a 1 μl syringe (Hamilton). The medium was changed and collected every two days. 6 days after GL261-EGFP inoculation, the slices were washed 3 times with PBS and fixed with 4% PFA at room temperature for 20min. Samples were washed with PBS and kept in 4℃ until further processing.

### CCK-8 kit assay

A total of 5,000 GL261-EGFP cells per well were seeded in a 96-well plate. Sixteen hours after plating, medium was change to DMEM only, or DMEM mixed with WT primary microglia conditional medium in a one-to-one ratio, or DMEM mixed with KO primary microglia conditional medium in a one to one ratio. The culture was kept in 37℃ incubator for 48 hours. CCK-8 reagent (Beyotime Biotechnology) was added (10 μl per well) and incubated for 2h. Plates were measured with a multireader at 450nm absorbance. Results were normalized to the absorbance of the control group DMEM only.

### Genotyping

Littermate controls were used in all experiments. For genotyping, genomic DNA was purified from tail biopsies by isopropanol precipitation, and the TREM2 locus harboring the knock-out mutation was amplified by polymerase chain reaction (PCR) using forward primer1 5’-CCCTAGGAATTCCTGGATTCTCCC-3’, forward primer2 5’-TTACACAAGACTGGAGCCCTGAGGA-3’and reverse primer 5’- TCTGACCACAGGTGTTCCCG-3’. PCR products were subsequently analyzed by 2% PCR Agarose gel electrophoresis. The band of wild type was detected at 231 bp and deletion in homozygous was detected at 316 bp.

### Immunofluorescence staining and image processing

Twenty-one days post-injection, mice brains were harvested and perfused with PBS followed by 4% PFA solution (Beyotime Biotechnology); 40 μm free-floating tissue sections were prepared as described above. Slices were washed three times with PBS for 5 min and blocked with 5% of donkey serum or 5% of goat serum in PBS with 0.5% Triton-X (93443, Sigma, Darmstadt, Germany). Primary antibodies were added overnight at 1:100 for CD31 (AF3628, R&D Systems, Minneapolis, USA), 1:500 dilution for Iba1 (ab5076, Abcam Cambridge UK or 019-19741, Wako, Tokyo, Japan), 1:400 for Ki67 (9129, Cell Signaling Technology, Danvers, USA), 1:500 for CD 8 (ab217344, Abcam) at 4° C. Iba1 and CD31 were detected using the secondary antibody Cy3-conjugated donkey anti-goat IgG (1:500, Beyotime), while CD8 and Ki67 were detected by Alexa-647-conjugated donkey anti-rabbit secondary antibody (1:500, Thermofisher). All brain slices were stained with DAPI(Beyotime) to detect the nuclei, and then were mounted in Aquapolymount (Polysciences, Warrington, USA).

Fluorescence images were taken with the apotome microscope or confocal microscope (Carl Zeiss) through Z-stack scanning under 20×objectives or 40×objectives, and the whole brain images were taken with TissueFAXS Q+Systems (TissueGnostics, Wien, Austria) with 20×objectives. 2-3 sections per biological replicate was analyzed and quantified for each mouse model, and the number of biological replicates per mouse model is indicated in the figure legends.

### Immunohistochemistry of human tissue material

Immunofluorescent staining was performed in human glioma tissues. The present study was approved by the Tongji Hospital Committee for human studies (process number:TJ-IRB20211162). Primary antibodies included Iba1 (1:500, Abcam), Trem2 (1:400, Cell Signaling Technology), and CD31 (1:800, Cell Signaling Technology). Briefly, tumor sections were deparaffinized, rehydrated, and antigen-retrieved by standard procedures. Then, samples were blocked with a PBS solution containing 1% BSA plus 0.3% Triton X-100 for 2 h at room temperature and then incubated with indicated primary antibody overnight at 4 °C followed by the fluorescence-conjugated second antibody (1:100, Proteintech, cat:SA00013-6, SA00009-2 and SA00013-1) at room temperature for 2 h. After being counterstained with DAPI for 5 min, sections were mounted on glass and subjected to microscopy. The images were acquired with a fluorescence microscope (Olympus (Tokyo, Japan), CKX53).

### RNA sequencing

GBM tissue and tools were pre-chilled with liquid nitrogen and the tissue was grinded into powder. A portion of the powder was transferred to TRIzol (Sigma) lysis buffer. RNA was extracted using a standard protocol (Guneykaya et al., 2018). Briefly, TRIzol lysis were mixed with 300 µl chloroform/isoamyl alcohol (24:1) and 2/3 volume of isopropyl alcohol was added. It was centrifuged at 17500×g for 25 min at 4°C. The supernatant was discarded, and the precipitation subsequently washed with 0.9 mL 75% ethanol and dissolved in 20 μl DEPC-H2O or RNase-free water. RNA quality was controlled using Agilent rna 6000 nano kit (5067-1511, Agilent, Santa Clara, USA) according to the manufacturer’s instructions. RNA was reverse transcripted into a cDNA library. Single-stranded circle DNA molecules are replicated via rolling cycle amplification, and a DNA nanoball (DNB) which contain multiple copies of DNA was generated. Sufficient quality DNBs are then loaded into patterned nanoarrays using high-intensity DNA nanochip technique and sequenced through combinatorial Probe-Anchor Synthesis (cPAS).

### Statistical analysis

The results are presented as mean ± standard mean error. Differences between groups were evaluated for significance by two-tailed Student’s test (two groups) or two-way analysis of variance (ANOVA) test (more than 2 groups) using GraphPad Prism 7 (GraphPad Software, San Diego, CA). Probability (P) values < 0.05 were considered statistically significant.

## Results

### The tumor volume is reduced in TREM2-deficient GL261 glioma male mice

To study the impact of TREM2 expression on glioma growth, we employed an *in vivo* mouse experimental glioma model. We injected GL261 mouse glioma cells expressing the fluorescent marker EGFP into the striatum of wild-type (WT) or TREM2 homozygous knock-out (KO) animals. Twenty-one days after tumor inoculation, mice were sacrificed and analyzed for tumor volume size and cellular parameters. Male and female animals were separately analyzed due to the fact that glioblastoma is more frequent in men than in women in a 1.6:1 ratio (Davis, 2016; Ostrom et al., 2018b). The tumor was visible from the surface of the cortex (Fig. 1A). The volume of the tumor was 4.7-fold larger in WT male than in WT female animals (47.86 ± 9.76 vs. 10. 27 ± 2.22 mm^3^, P = 0.0001) (Fig.1 B & C). In the female group, TREM2 deficiency did not show a significant impact on tumor size (WT 10.27 ± 2.22 vs. KO 10. 07 ± 2.84 mm^3^) (Fig.1 B & C). In contrast, in the male animals, the tumor volume was significantly smaller in KO animals as compared to the WT counterparts (WT 47.86 ± 9.76 vs. KO 16.73 ± 4.72 mm^3^, P = 0.0018) (Fig.1 B & C). Interestingly, tumor volume in TREM2- deficient male animals shows no differences compared to female WT or female KO animals, suggesting the gender effects for GBM could potentially be attributed to TREM2 expression. Collectively, these findings suggest that the GL261 GBM mouse model recapitulate the gender preference, most importantly TREM2 expression promotes tumor progression in male animals. Therefore, we used only the male GL261 GBM mouse model for the subsequent analysis of the mechanistic impact of TREM2 expression on glioma progression.

**Figure 1.**
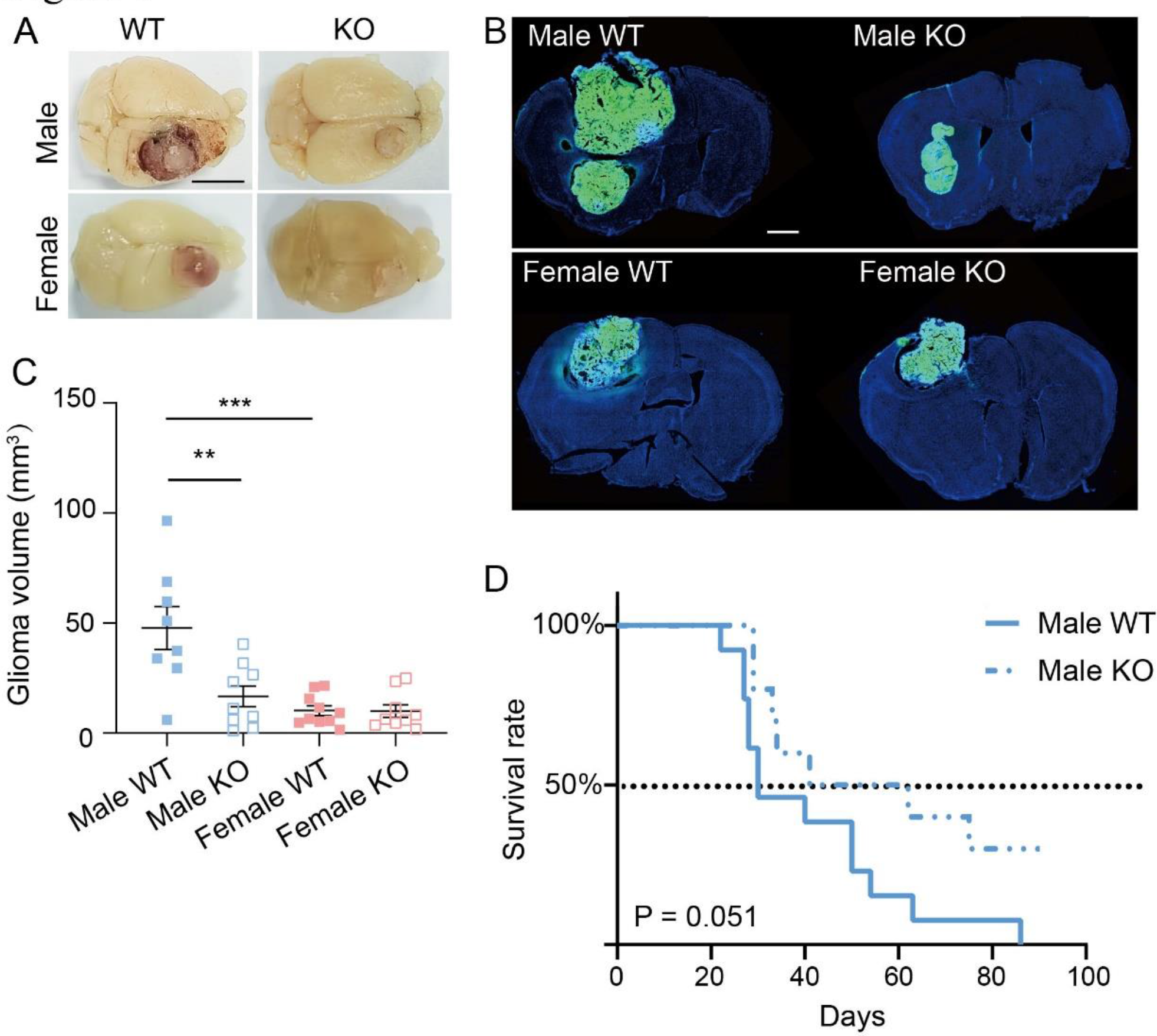
The tumor volume is reduced in TREM2-deficient GL261 glioma male mice **A.** Representative images of whole brains with tumors of male and female WT and TREM2 KO mice. Scale bar denotes 1 cm. **B.** Representative immunofluorescence images of WT and TREM2 KO coronal brain sections 21 days after stereotactic injection of GL261-EGFP into male and female mice. GL261-EGFP (green); DAPI (blue) (Scale bar, 1 mm). **C.** Quantification of glioma volume based on the EGFP signal. Female WT (n=10), Female KO (n=9); Male WT (n=8), Male KO (n=9); Data are presented as means ± SEM. One-way ANOVA with Bonferroni corrected multiple comparisons. **P < 0.01; ***P < 0.001. **D.** Kaplan–Meier curves represent the cumulative survival of male WT and TREM2 KO mice injected with GL261-EGFP cells (n= 10-13 per group, Log-rank test P=0.0512).

We further tested the impact of TREM2 on the survival time of tumor-inoculated male mice. Mice were scored daily after 21 days for the following symptoms: body weight, hunchback, low reaction to stimulation, tarnished fur, or movement problems. Animals were sacrificed if they show severe symptoms in the criteria mentioned above. The difference on survival probability between these two genotypes was non-significantly different when 90 days were assigned as observation endpoint. (median survival: WT 30 days vs KO 51.5 days, P = 0.051) (Fig.1 D).

### TREM2 deficiency results in lower GAMs density at the tumor core and rim in male glioma animals

Since TREM2 is expressed by microglia as well as by infiltrating macrophages, we determined the distribution of GAMs in different regions of the GL261-EGFP inoculated animals. We determined the density of GAMs at the core of the tumor, at the outer rim of the tumor, in the area of adjacent brain tissue denoted as interface (0-0.3 mm away from the EGFP positive cells). Moreover, we analyzed the density in four regions with increasing distance (in 0.3 mm steps) from the tumor (adjacent brain region ABR 1, 0.3 - 0.6 mm; ABR 2, 0.6 - 0.9 mm; ABR 3, 0.9 – 1.2 mm; ABR 4, 1.2 – 1.5 mm) (Fig. 2 A & B & C) and at the contralateral side of tumor inoculated hemisphere. In the core of the tumor, the coverage of the Iba1-fluorescent area is significantly higher in WT compared to TREM2 KO animals (12.46% ± 1.27% vs. 6.91 ± 1.00, P=0.0002) (Fig. 2 B & D). Similarly, WT animals also show higher Iba1- fluorescent area at the rim of tumor (10.75% ± 0.95%, vs. 6.02 ± 0.88, P=0.0024) (Fig. 2 C & D). However, the Iba1-fluorescent area is not significantly different between the two genotypes in the tumor-brain interface, the ABR1-4 (Fig. 2 C & D) and the contralateral area (Fig. 2 B & D). Notably, the distribution of GAMs at the tumor-brain interface is denser compared to the rim of the tumor (Fig. 2 C & D), similar to the data that have been reported in patient glioblastoma tissue (Yin et al., 2022), indicating that the GL261 glioblastoma mouse model represents the clinical features of GAMs distribution.

**Figure 2.**
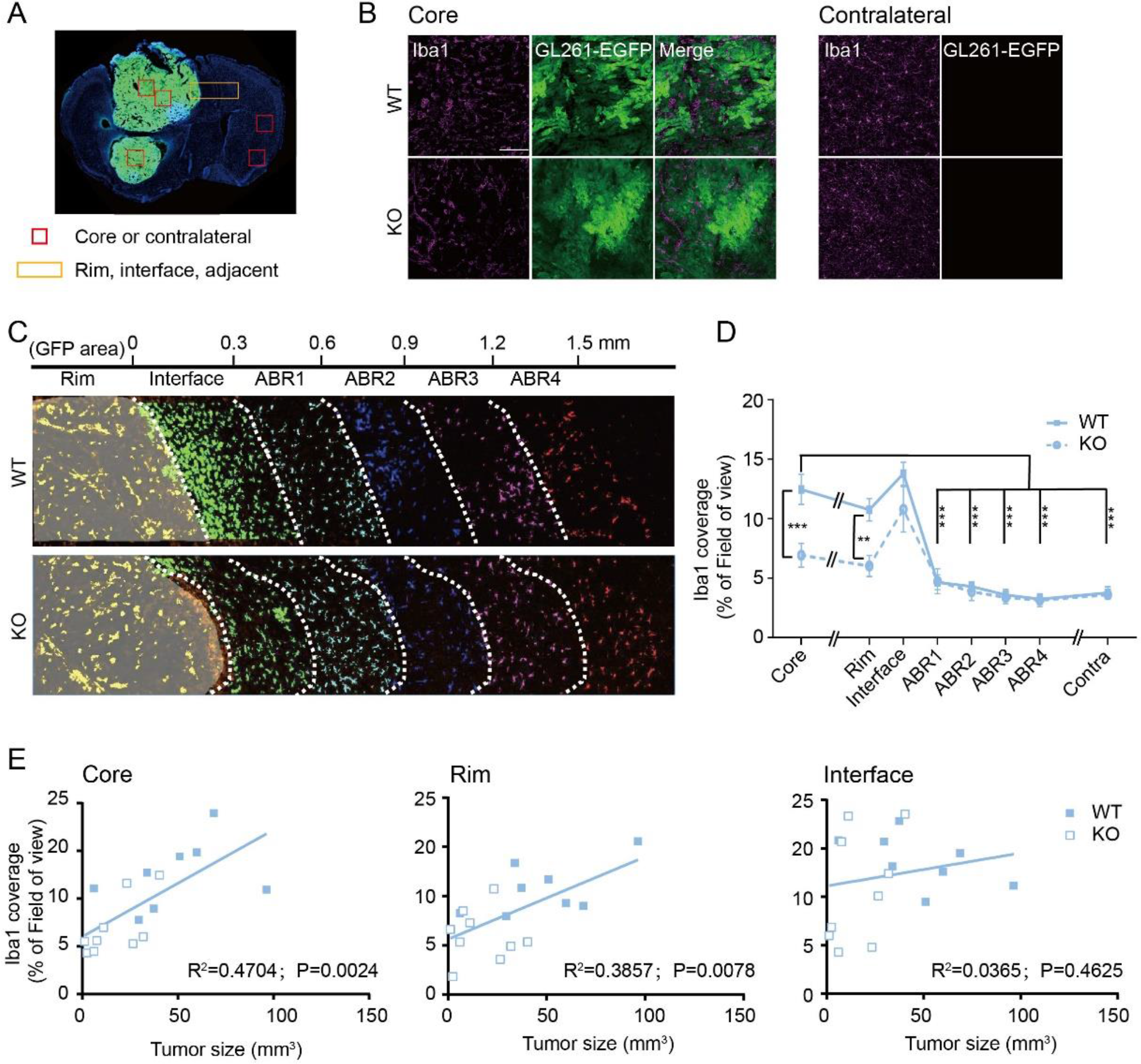
TREM2 deficiency results in lower GAMs density at the tumor core and rim in male glioma animals **A.** Overview image showing the regions analyzed for coverage of microglia/macrophages in a tumor bearing male mouse brain. The core region of glioma and contralateral region were chosen with a 0.5×0.5mm box and the rim region was chosen with 0.5×2.5mm box. **B.** Representative images of Iba1^+^ cells in the glioma core and contralateral hemisphere from WT (top) and TREM2 KO (bottom) male mice. Iba1 (magenta), GL261-EGFP (green), and merged image. Scale bar denotes 100 μm. **C.** Representative immunofluorescence images for microglia/macrophages (Iba1), glioma cells (GL261-EGFP) and merged image of WT and TREM2 KO mice at the rim of glioma including glioma area (up to 0 mm), glioma interface (0-0.3 mm) and adjacent brain region (0.3-1.5 mm). Different color codes for Iba1^+^ cells in different sub divided regions. **D.** Quantification of the percentage of the Iba1 coverage in the analyzed regions as described in C (WT n=8, KO n=9). Data are presented as means ± SEM. *P < 0.05,**P < 0.01,two-way ANOVA with Bonferroni corrected multiple comparisons. **E.** Correlation between Iba1 coverage and glioma size in the core (left), rim (middle) and the interface (right) as defined in C.

We also tested the correlation between the density of GAMs and glioma volume. GAMs density is positively correlated with glioma volume in the core and rim region of glioma (Fig. 2 E left and middle panel). However, there is no positive correlation between the glioma volume and the density of GAMs in the interface region (Fig. 2 E right panel). These data suggest that the density of GAMs within the glioma correlates with glioma size. More importantly, these data suggest the reduced GAMs density in the core and rim of tumor in the TREM2 KO animals correlates with a smaller tumor size.

To further characterize the immune environment of the glioma tissue, we analyzed the density of CD8^+^ T cells within the tumor. The density of CD8^+^ T cell population is not influenced by the TREM2 expression (Supplement fig. 1 A & B).

### TREM2 deficiency leads to a lower proliferation rate in glioma cells, **but not in GAMs**

We next attempted to study the proliferation of glioma cells, GAMs and remaining cell types in the tumor environment determined by Ki67 immunofluorescence staining. These experiments were performed in male mice only. The Ki67 signal was colocalized with DAPI. The Ki67^+^ cell number was normalized to total DAPI^+^ cell number in a given region. We found that the percentage of Ki67 positive cells is lower in TREM2 KO GBM as compared to WT (Fig. 3 A & B). We further divided the Ki67^+^ cells into three categories, namely in Iba1^-^ / EGFP^+^ (glioma cells), Iba1^+^ / EGFP^-^ (GAMs), Iba1^-^/ EGFP^-^ (other brain cells) (Fig. 3 C & D). In the GL261 tumor cell population, Ki67 is significantly higher in WT animals compared to TREM2 KO animals, indicating that the proliferation rate of tumor cells is reduced due to the loss of TREM2 (Fig. 3 C & D). In the GAMs population, the Ki67^+^ population is not significantly different in WT versus KO animals (Fig. 3 C & D), suggesting that TREM2 deficiency did not show a significant impact on the proliferation rate of GAMs. Similarly, we found no significant difference in Ki67^+^ cells in WT and KO animals in the population of Iba1^-^ / EGFP^-^ (other brain cells) (Fig. 3 C & D).

**Figure 3.**
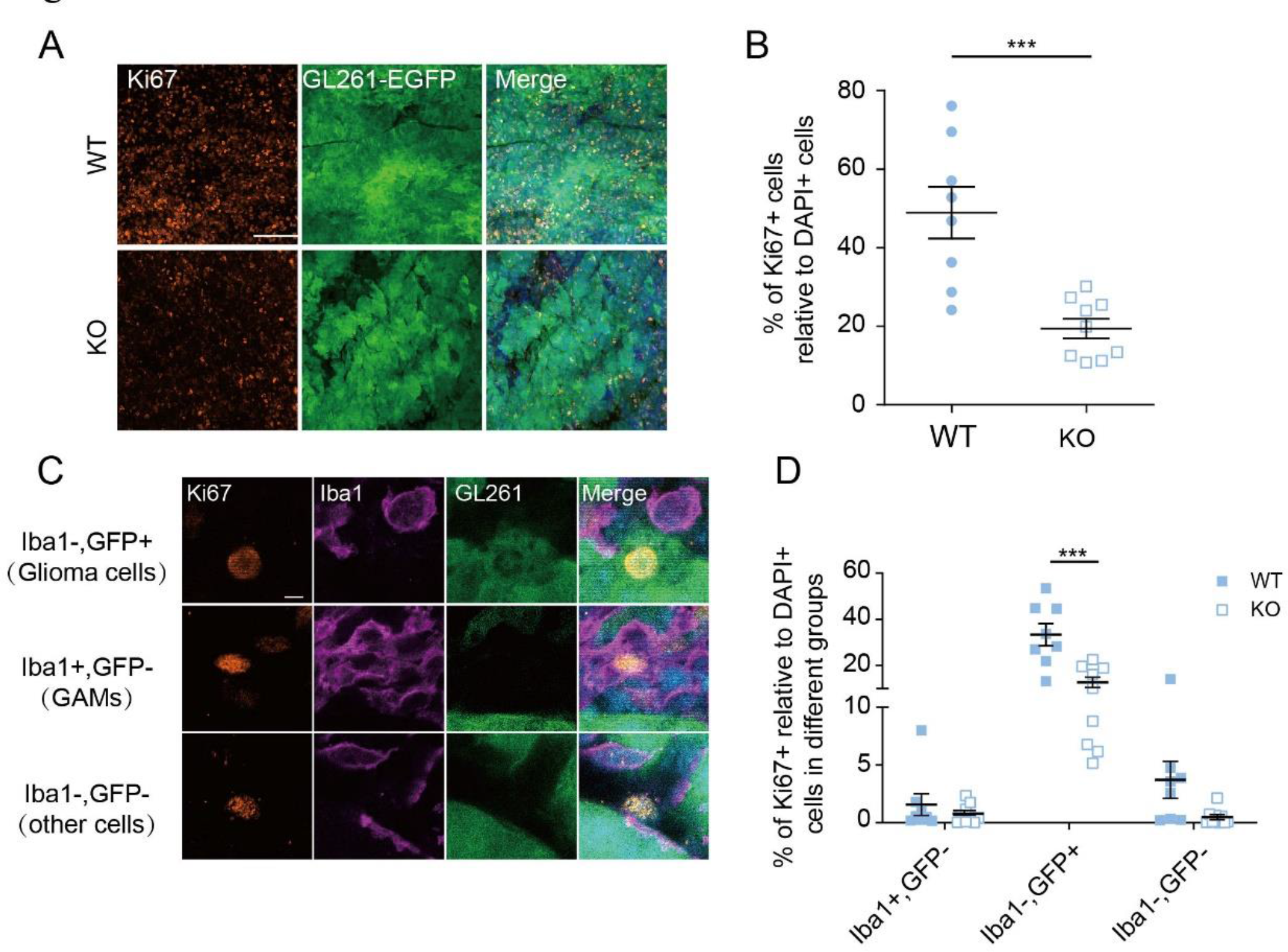
TREM2 deficiency leads to a lower proliferation rate in glioma cells, but not in GAMs **A.** Images of Ki67 staining (left), EGFP fluorescence of GL261 glioma cells (middle) and merged image (right) for WT (top) and TREM2 KO mice (bottom). The bar denotes 100 μm. **B.** Quantification of the percentage of Ki67^+^ cells among the DAPI^+^ cells in the glioma tissue of WT and TREM2 KO male mice. **C.** Corresponding images of Ki67, Iba1, EGFP (from GL261-EGFP cells) and merged labelling showing examples of Ki67 cells co-labelled with glioma cells (top, Iba1-, EGFP+), GAMs (middle, Iba1+, EGFP-) and other cells (lower, Iba1-, EGFP-) in glioma. The bar denotes 5 μm. **D.** Quantification of the images shown in C. Data are presented as means ± SEM. Two- way ANOVA with Bonferroni corrected multiple comparisons. ***P < 0.001.

### The density of CD31^+^ blood vessels in the glioma is reduced in the TREM2-deficient mice

Since angiogenesis is an important factor in malignant gliomas influencing tumor growth and progression (Fischer et al., 2005), we quantified the density of CD31^+^ blood vessels in the tumor core, the tumor rim and adjacent brain regions, similar to the analysis for Iba1 coverage as described above (Fig.2A). As expected, the density of blood vessels (CD31^+^ coverage) is increased in the core, rim and tumor-brain interface region as compared to the contralateral side in both genotypes, indicating a strong angiogenesis in the tumor-related environment (Fig.4A & B & C). The density of CD31^+^ blood vessels in the areas further away from tumor (0.3-1.5 mm, defined as ABR1-4 similar as described above for microglia density analysis) is not significantly different compared to the contralateral site in both genotype (Fig.4 B & C). These results indicate that the increase in vessel density is restrict to the glioma tissue and its closely adjacent region (designated as interface). Importantly, in the glioma core, rim and the tumor-brain interface region, the coverage of CD31^+^ regions is higher in WT compared to TREM2 KO animals (Core: WT 9.159% ± 1.216% vs. KO 5.272 ± 1.246, P=0.0033; Rim: WT 11.878% ± 1.430% vs. KO 5.277 ± 1.075, P<0.0001; Interface: WT 6.629% ± 1.461% vs. KO 4.043 ± 0.808%, P=0.048) (Fig.4A & B & C). Next, we analyzed the vessel length in the core region. The CD31^+^ vessel length in the core region is significantly longer in the WT compared TREM2 KO animals (Fig 4 D).

**Figure 4.**
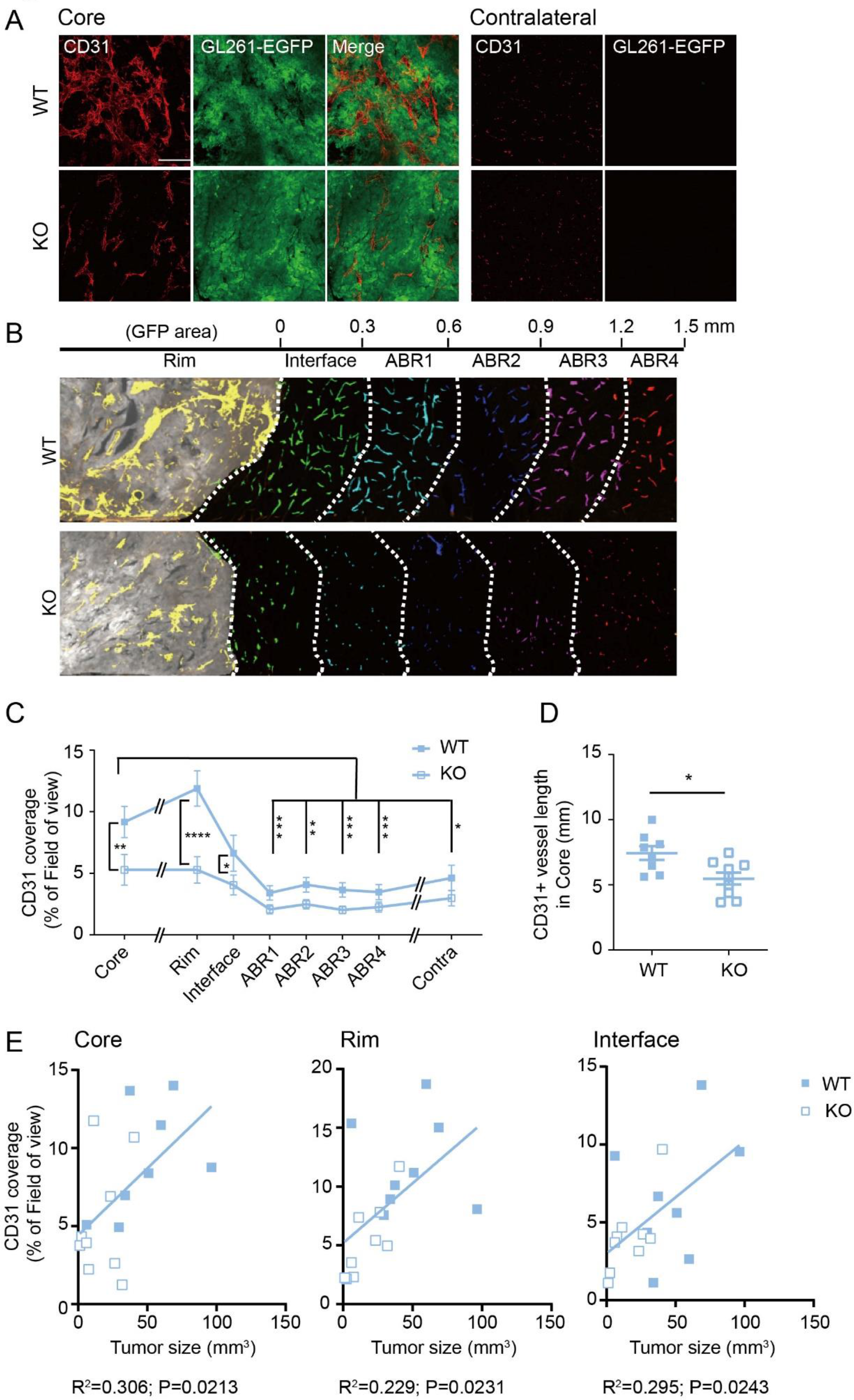
The density of CD31^+^ blood vessels in the glioma tissue is reduced in TREM2-deficient mice **A.** Representative images of vessels in the core of the glioma and contralateral site of WT (top image) and TREM2 KO (bottom image) male mice. Immunofluorescence staining for CD31 (magenta), EGFP (for GL261-EGFP cells) (green) and a merged image. Scale bar denotes 100 μm. **B.** Representative images for vessels (CD31^+^) of WT and TREM2 KO animals at the rim and interface of glioma. Different color codes for vessels in different sub divided regions as described in detail in the legend to Fig. 2. **C.** Quantification of the percentage of CD31^+^ immunofluorescence coverage in the regions as defined in B. (WT n=8, KO n=9) Data are presented as means ±SEM. *P < 0.05; **P < 0.01; *** P < 0.001; ****P < 0.0001, two-way ANOVA with Bonferroni corrected multiple comparisons. **D.** Quantification of vessel length based on CD31 staining in the glioma core of WT and TREM2 KO. **E.** Correlation between CD31 coverage and glioma size in the core (left), rim (middle) and the interface (right) as defined in the legend to Fig. 2.

The glioma size is positively correlated with CD31^+^ area in core, rim and interface regions, respectively (Fig. 4E), suggesting a positive role of vascularization on tumor growth. These data suggest that the TREM2 deficiency attenuates angiogenesis in the glioma tissue.

### TREM2 deficiency suppresses angiogenic genes and MHC clusters

To understand the underlying mechanisms how TREM2 deficiency lead to reduced vasculature, we performed a transcriptomic analysis of the glioma tissue. Twenty-one days after GL261-EGFP inoculation, the glioma tissue was dissected from WT or TREM2 KO male mice for RNA extraction and bulk RNA-seq. A similar number of genes was detected in both genotypes (16353 ± 207 in WT vs. 16208 ± 23 in KO). At the settings of log2|FC| ≥1 and adjusted p value <0.05, 144 downregulated and 3 upregulated genes were discovered (Fig. 5A and Supplementary fig. S2A). Samples from WT and TREM2 KO animals show a clear separated cluster, and 144 out of 147 differentially expressed genes (DEGs) are downregulated in TREM2 KO samples (Fig. 5 A & B, Supplementary fig. S2B). KEGG pathway analysis was carried out to identify involved signaling pathways for TREM2 functions in glioma pathology (Fig. 5 C). KEGG category analysis showed that a large proportion of DEGs were immune genes which may participate in immune diseases (28 genes), cancers (24 genes) and, signaling molecules and interaction (25 genes, signaling molecules and receptors) (Fig. 5 C). Interestingly, we noticed that a group of cell adhesion molecules in KEGG pathway analysis were downregulated in KO glioma, including eleven major histocompatibility complex (MHC) I and MHC II cluster genes (MHC I cluster: H2-D1, H2-M3, H2-Q4, H2-Q6, H2-Q9, H2-T22, H2-T23, H2-T24; MHC II cluster: H2-Ab1, H2-DMb1, H2- Eb1) (Fig. 5 C & D). The reduction of MHC gene cluster may suggest a reduced antigen presentation in GBM from KO animals compared to WT, and may contribute to immune escape and impaired functions of CD4^+^/CD8^+^ cells and NK cells in GBM (Burster et al., 2021; Kilian et al., 2023).

**Figure 5.**
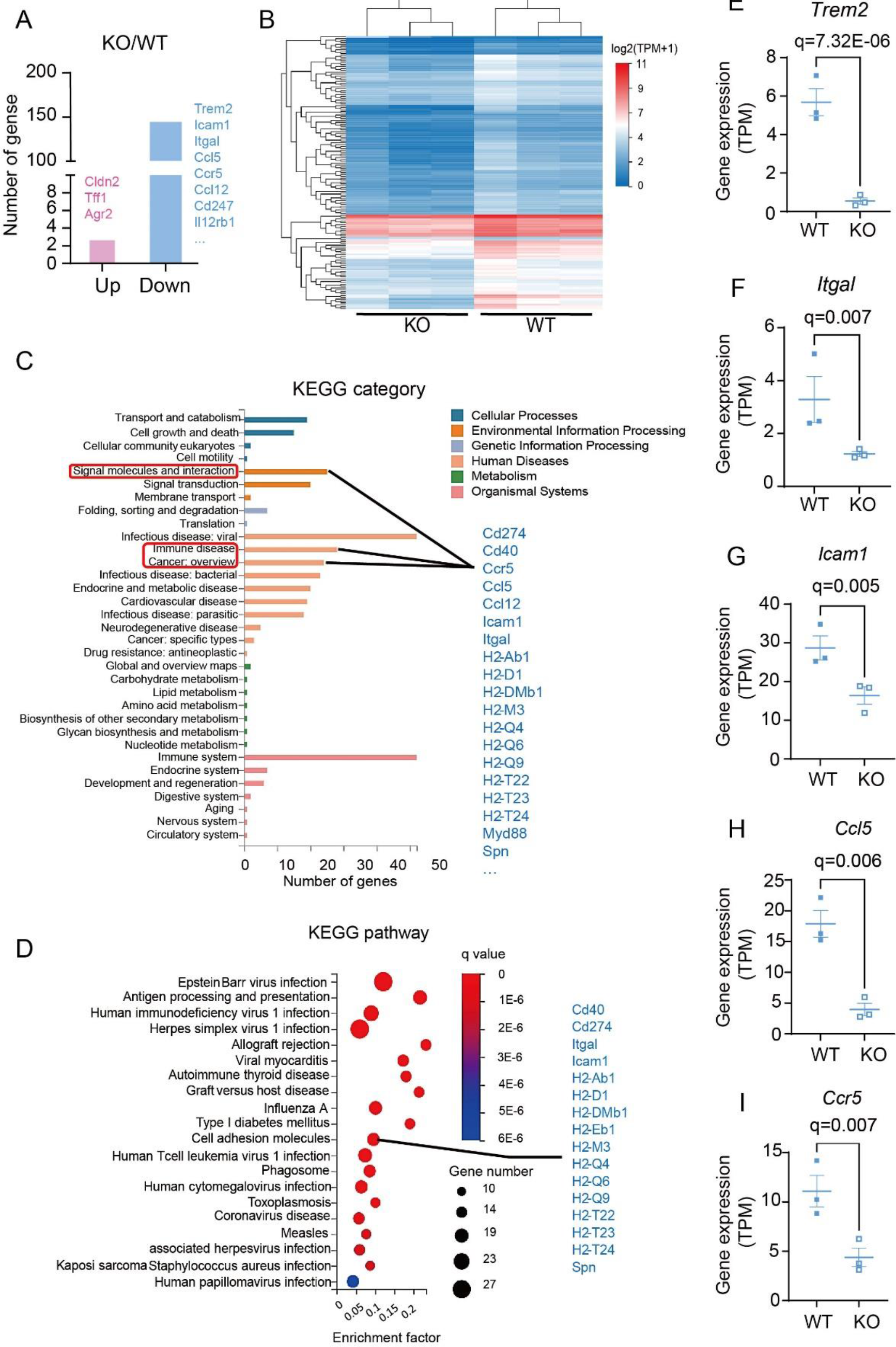
TREM2 deficiency suppresses angiogenic genes and MHC clusters **A.** The number of differentially expressed genes (DEGs) in the GBM sample in TREM2 KO mice compared to WT mice. 144 downregulated (some examples are indicated) and 3 upregulated genes were identified. **B.** Heat map of log2 transcripts per kilobase million +1 (log (TPM+1)) of 147 DEGs from WT and TREM2 KO samples with adjusted p value <0.05 and log2|FC| ≥1. **C-D.** KEGG category (C: genes listed on the right refers to the categories as indicated) and pathway (D) analysis of DEGs, the genes listed at the right refer to genes in Cell adhesion molecules pathway. **E-I.** Gene expression (TPM) of *Trem2* (E), *Itgal* (F), *Icam1* (G), *Ccl5* (H) and *Ccr5* (I) (N=3). q values (adjusted p value) are indicated.

Some pro-angiogenic genes or immune signaling genes which can modulate angiogenesis were also identified in the downregulated group (KEGG pathway category: signaling molecules and interaction) (Fig. 5 C), including: two cell adhesion molecules Icam1 (Lim et al., 2022) and Itgal (Silva et al., 2008; Zhang et al., 2022a)), chemokines Ccl12 and Ccl5 (Suffee et al., 2011; Wang et al., 2015), the chemokine receptor Ccr5 and inflammatory signaling adaptor Myd88 (Zhang et al., 2020). Based on the adjusted p value (q value), these genes are downregulated in GBM from TREM2 KO animals (Figure 5E-I, Supplementary fig. S2C and S2D). In contrast, one of the three upregulated genes in KO animals (Fig. 5 A and Supplementary fig. S2E), Tff1 (Shi et al., 2019; Zhang et al., 2022b), inhibits angiogenesis in cancer progression, which support the notion that TREM2 positively regulates angiogenesis in glioma environment.

In addition, we examined whether TREM2 correlated with these pro-angiogenic or anti-angiogenic genes in glioma patients. In datasets from Chinese Glioma Genome Atlas (CGGA) (Zhao et al., 2021) RNA-seq dataset, TREM2 expression was found to be positively correlated with Itgal, Icam1, Ccl5, Ccr5, and Myd88 expressions in glioma patients (Supplementary fig. S43A-E). Tff1 expression can only be found in one of the three microarray datasets and was negatively correlated to TREM2 expression in recurrent glioma patients (Supplementary fig. S3F).

In summary, the transcriptomic analysis revealed that TREM2 depletion on one hand inhibits immune genes, in particular MHC I and II clusters, which may suppress antigen presentation. On the other hand, TREM2 depletion leads to decreased pro-angiogenic and increased anti-angiogenic gene expression.

### TREM2 shows no impact on glioma growth in isolated *ex vivo* and *in vitro* systems

To study if a functional vasculature system is a prerequisite for TREM2-mediated impact on glioma size, we analyzed glioma growth in the organotypic brain slice glioma model (OBS) where the tissue contains all the major brain cells including microglia, but lacks a functional vasculature system and an influence on the peripheral immune system including the immigration of blood monocytes. Slices were prepared from postnatal day 5-8 WT or TREM2 KO animals, cultivated for 5 days and subsequently injected with 5000 GL261-EGFP cells (Fig.6 A). Six days after glioma cell inoculation, the volume occupied by EGFP-labeled glioma cells was determined by immunohistochemistry and confocal microscopy with z stack images. The volume of glioma is not significantly different between WT and KO, indicating that TREM2 has no effect on glioma growth in the isolated *ex vivo* model (Fig.6 B & C). In addition, no difference was found regarding the density of Iba1^+^ cells in the glioma area of WT and KO OBS tissue (Fig.6 D & E). TREM2 is expressed by Iba1^+^ microglia, as shown by colocalization of TREM2 and Iba1 in the WT OBS (Fig.6 D), and the specificity of the immunostaining is shown by lack of TREM2 signal in the KO tissue (Fig. 6 D). After normalized to Iba1^+^ area, TREM2 coverage is higher within glioma compared to tumor free tissue (Fig.6 F & G), suggesting that glioma inoculation stimulates TREM2 expression in WT OBS. As expected, no CD31^+^ vessel can be detected within the glioma area in this *ex vivo* model (Fig. 6H) To further validate if TREM2 deficiency in microglia would have a direct impact on glioma cell proliferation, we incubated GL261 cells with conditional medium from WT or TREM2 KO primary microglia, followed by Cell Counting Kit-8 (CCK-8) assay to quantify cell density and viability. In line with previous reports (Huang et al., 2020), microglia conditioned medium strongly stimulated the proliferation of GL261 cells resulting in higher viability as compared to unstimulated cells (Supplementary fig. S4 A). However, we found no difference of viable glioma cell numbers when stimulated with either WT or TREM2 KO microglia-produced conditioned medium (Supplementary fig. S4A), supporting the view that TREM2 deficient microglia do not affect glioma growth via soluble factors.

**Figure 6.**
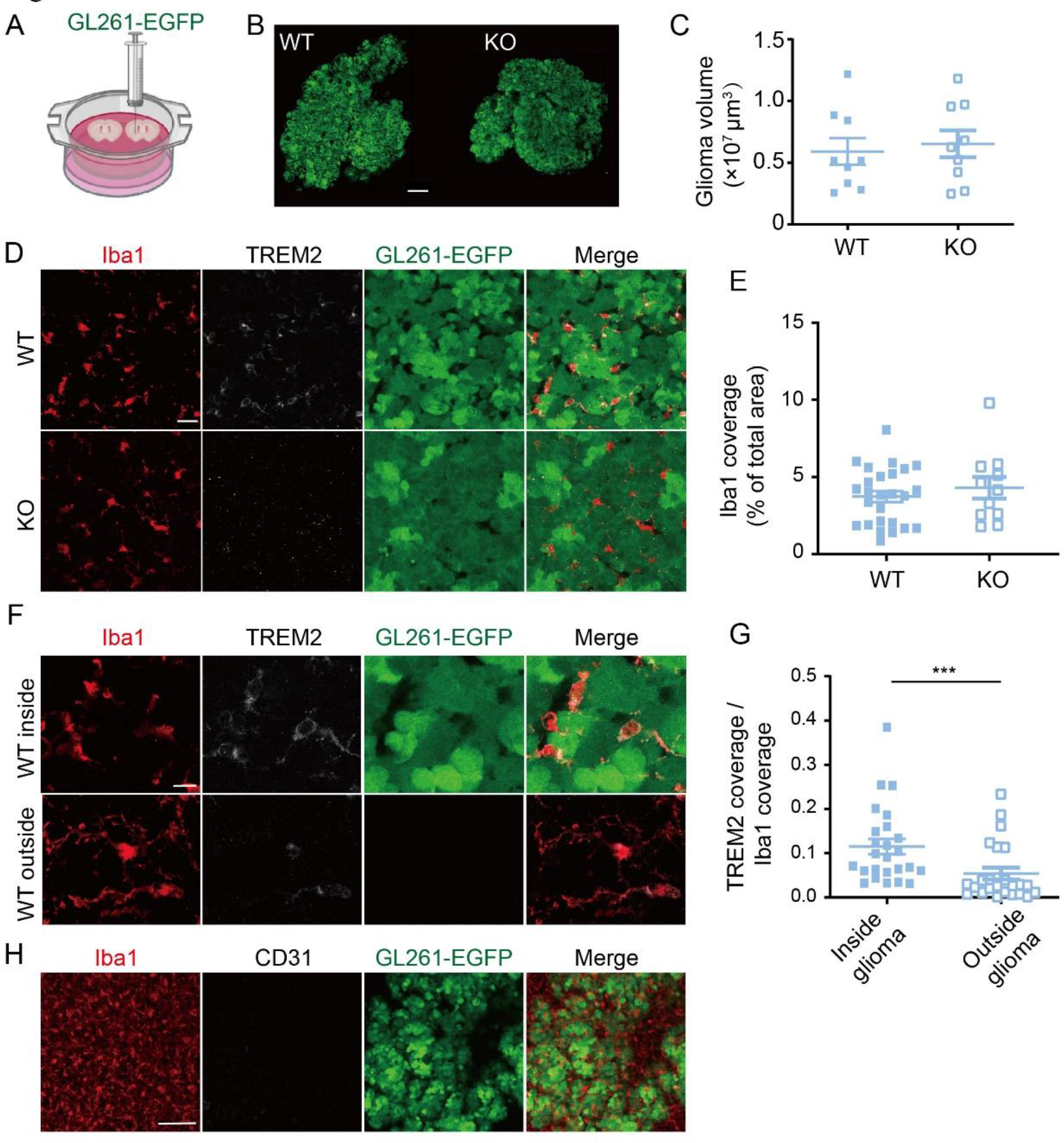
TREM2 show no impact on glioma growth in isolated *ex vivo* system **A.** Schematic view of the organotypic brain slice culture (OBS) inoculated with GL261- EGFP glioma cells. **B.** Representative immunofluorescence images of glioma (green) in WT and TREM2 KO OBS. (Scale bar, 200 μm) **C**. Quantification of glioma volume based on the EGFP signal. **D.** Representative image for Iba1 (red), TREM2 (white), GL261-EGFP (green) and merge image of WT (top) and TREM2 KO (bottom) from OBS glioma model. (Scale bar, 20μm) **E.** Quantification of the percentage of Iba1^+^ covered area in the glioma region (GL261-EGFP). **F.** Representative image for Iba1(red), TREM2 (white), GL261-EGFP (green) and merged image of a region inside (top) and outside (bottom) of the glioma in a slice from a WT animal. (Scale bar, 10 μm) **G.** Quantification of the percentage of TREM2^+^ covered area (normalized to Iba1^+^ area) inside and outside of glioma. **H.** Representative immunofluorescence image of Iba1 (red) and CD31 (white) and EGFP (green) within the glioma region in WT and TREM2 KO OBS. (Scale bar, 100μm). Data are presented as means ± SEM. ***P < 0.001, student t-test.

### TREM2 is upregulated in GAMs in human glioma tissue and negatively correlates with patient survival

To evaluate the TREM2 expression level in the human glioma environment, we immunostained human brain sections from eleven GBM and two epilepsy patients for TREM2 and Iba1, as a marker for GAMs. The TREM2 level is higher in GBM cases compared to epilepsy tissue (Fig. 7 A & B). Most importantly, most of the TREM2 positive signal is co-localized with Iba1 in the GBM cases (Fig. 7 A & B), suggesting TREM2 is expressed by GAMs in the tumor environment.

**Figure 7.**
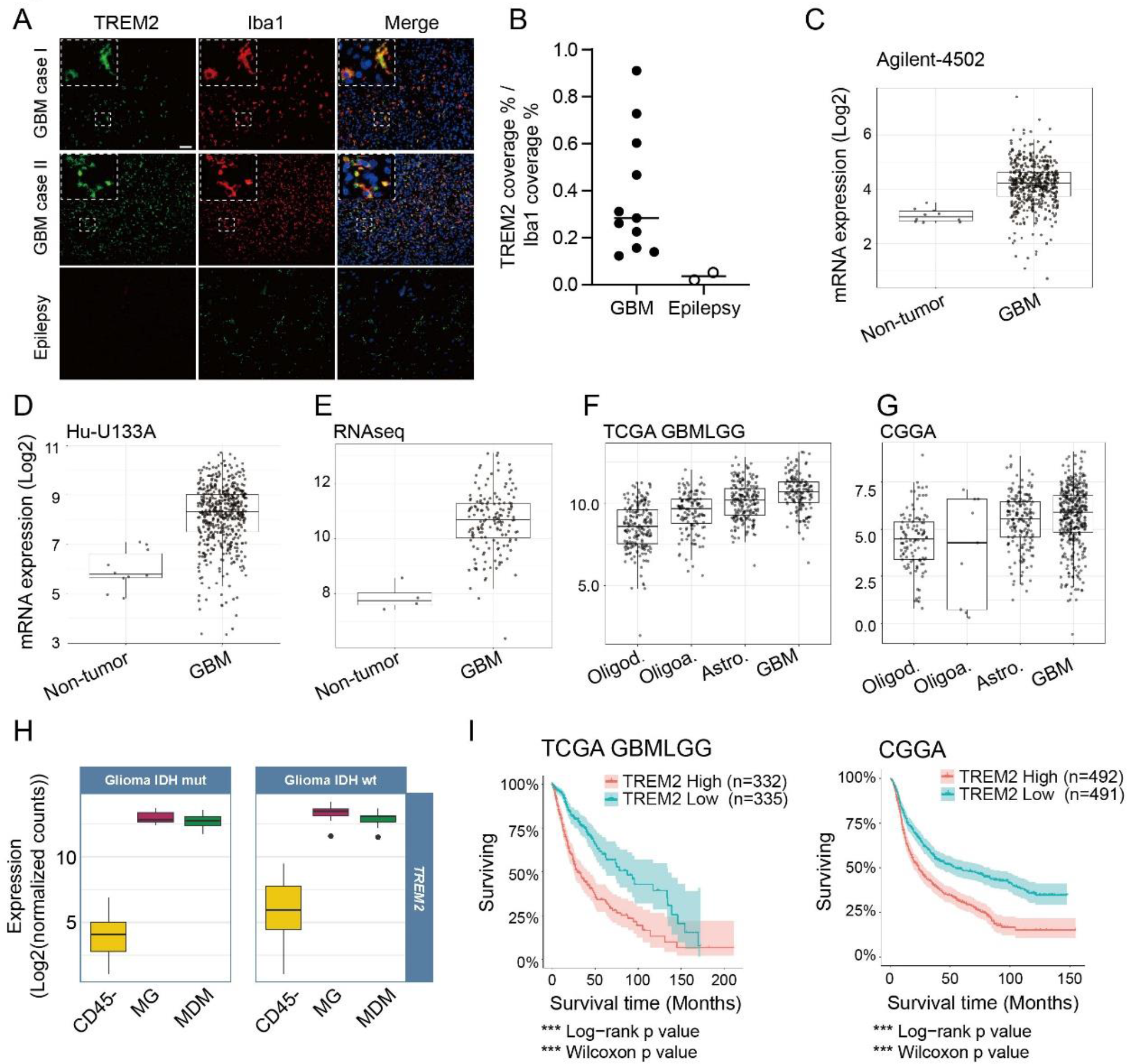
TREM2 is upregulated in GAMs in human glioma tissue and negatively correlates with patient survival **A.** Representative images of TREM2 (green), Iba1 (red) and merged images from two GBM patient samples (top and middle) and epilepsy (bottom) tissue from a patient. The insets at the top left are four times enlarged images of the region delineated by the square in the center of the image (Scale bar, 50 μm) **B.** Corresponding quantification of TREM2^+^ area (normalized to Iba1^+^ area) from eleven GBM samples and two epilepsy samples. **C-G.** TREM2 mRNA expression level from different datasets: TCGA- Agilent-4502 (C), TCGA-Hu-U133A (D), TCGA-RNAseq (E), TCGA-GBMLGG (F), CGGA (G). In C, D, E GBM tissue is compared to non-tumor tissue, while in E and F oligodendro-glioma, Oligo-astrocytoma, astrocytoma and GBM cases are compared. **H.** TREM2 mRNA expression level in different cell types (CD45-negative cells, MG: microglia; MDM: monocyte derived macrophages) based on the Brain Tumor Immune Micro Environment database distinguishing GBM with IDH mutations and IDH wildtype. **I.** Survival probabilities of patients with high and low TREM2 expression level. Left: dataset from TCGA GBMLGG; Right: dataset from CGGA.

To confirm the evaluated TREM2 expression in the human glioma environment, we first performed a systematic meta-analysis on The Cancer Genome Atlas Program (TCGA) GBM-Agilent 4502A (Fig.7 C), TCGA GBM-HU133A (Fig.7 D), TCGA GBM-RNA-seq (Fig.7 E), TCGA GBMLGG RNA-seq (Fig.7 F), CGGA (Fig.7 G) database utilizing the bioinformatics platform GlioVis (http://gliovis.bioinfo.cnio.es/) (Bowman et al., 2017). The expression of TREM2 is higher in GBM compared to non-tumor samples in all datasets (Fig7. C-E). Moreover, TREM2 expression is higher in GBM compared to other types of glioma in both TCGA GBMLGG and CGGA datasets (Fig.7 F & G). TREM2 is expressed by cells from the myeloid linage, including microglia, macrophages and dendritic cells (Colonna, 2003; Xiang et al., 2016). To further study the expression pattern of TREM2 in GBM, we analyzed the Brain Tumor Immune Micro Environment dataset (https://joycelab.shinyapps.io/braintime/) (Klemm et al., 2020). The expression of TREM2 is significantly higher in microglia and monocyte derived-macrophages (MDM) compared to CD45- cells, and this enrichment of TREM2 in microglia and MDM is regardless of the isocitrate dehydrogenase (IDH) genetic status (Fig. 7 H). Our immunohistochemistry analysis of tissue from GBM patients is in line with these datasets, suggesting that TREM2 is highly expressed by GAMs in the GBM environment.

To assess the correlation of TREM2 expression and the probability of survival in glioma patients, we again used the GlioVis data portal for analysis (Bowman et al., 2017). In two different datasets, TCGA GBMLGG and CGGA, patients with higher TREM2 expression show significantly lower survival probability (Fig. 7 I), indicating high TREM2 expression is detrimental for glioma patients, which is similar to what we detected in the GL261 mouse glioma model.

## Discussion

In many neurodegenerative disorders, including Alzheimer’s disease (Lewcock et al., 2020; Schlepckow et al., 2020; Wang et al., 2020), amyotrophic lateral sclerosis (Xie et al., 2022), frontotemporal dementia (Kleinberger et al., 2017), TREM2 has a beneficial role. In contrast, in the MCA/1956 sarcoma model, TREM2 has a detrimental role on tumor growth by modulating the immune landscape (Katzenelenbogen et al., 2020; Molgora et al., 2020). Although it has been shown that TREM2 is highly expressed in the glioma environment (Molgora et al., 2020), the functional role of TREM2 in glioma is largely unknown. In the current study, we used the GL261 glioma mouse model to show that TREM2 deficiency leads to a reduced glioma size, results in less accumulation of GAMs in the glioma tissue and an attenuated vessel growth. TREM2 acts in a gender-specific fashion in that these TREM2-dependent changes were only observed in male, but not in female animals. This uncovered a previously unknown gender-dependent feature of TREM2 function.

This is in line with patient data which show a clear sex bias in the incidence of glioma, namely a male to female ratio of glioma patients of about 1.6 to 1 (Ho et al., 2014; Ostrom et al., 2018a). Moreover, female glioma patients have a longer survival time than male (Ostrom et al., 2018b). It has even been suggested that therapeutic strategies should be optimized depending on treatment in a sex-specific manner (Massey et al., 2021). Moreover, our data indicate that the male mice had a larger tumor volume compared to female mice, suggesting that this model can be exploited to study the gender impact on glioma growth.

In the last years it has become apparent that microglia from male and female mice are quite distinct (VanRyzin et al., 2018, 2020). Microglia in male and female mice differ in structure, function, and transcriptomic and proteomic profiles. Male microglia are more frequent in specific brain areas, have a higher antigen-presenting capacity, and appear to have a higher potential to respond to stimuli such as ATP (Guneykaya et al., 2018). Sex-specific differences in microglial functions and features must not necessarily be due to intrinsic genetic differences but could also be influenced by extrinsic sex-specific cues. Deletion of neuroligin-4, a protein expressed by neurons and not by microglia, is linked to autism spectrum disorders and has a sex-specific impact on microglia. Male microglia show a lower cell density, a less ramified morphology, a reduced response to injury and purinergic signaling specifically in the hippocampal CA3 region (Guneykaya et al., 2023).

An important issue is how a microglia-specific gene deletion, namely of TREM2, affects glioma growth and angiogenesis in combination with a lower number of microglia in the tumor. What are the pathways of that interaction? In our in vitro studies we applied supernatant from WT and TREM2 KO microglia to glioma cells and determined glioma cells number. While we confirmed former studies that the microglia supernatant increased glioma cell number, there was no significant difference between WT and TREM2 KO. This implies that soluble factors derived from microglia cannot explain the effect of TREM2 KO on glioma growth (with all the cautions relying on an in vitro model). We cannot exclude that physical contact might be a factor, but it is more likely that the glioma cell growth is indirectly affected by the TREM2 deletion.

Blood vessels in the tumor are larger and more frequent than in the normal tissue with irregular lumen diameters, they are dilated and highly permeable, and branch irregularly (De Bock et al., 2011). Angiogenesis is essential for tumor growth and progression (De Bock et al., 2011). As expected, we found that the vessel density is significantly higher within the glioma as well as in the glioma brain interface region. More importantly, the tumor size is positively correlated with vessel density in the glioma region. We found that the density of CD31^+^ vessels is significantly higher in glioma region from WT animals compared to TREM2 KO. This could lead to the hypothesis that microglial TREM2 indirectly promotes glioma progression via enhancing the angiogenesis in the glioma environment. We also found that the total vessel length is longer in the core region of glioma from WT animal compared to KO. Interestingly, TREM2 deletion shows no impact on glioma size in the isolated OBS glioma model where no functional vasculature system is present, arguing a TREM2-dependent angiogenic effect could promote glioma progression. It should, in addition, be noted that the OBS glioma does not only lack a functional vasculature system, but also lacks the immigrating cells from the blood system such as monocytes. Both features could play a role.

One possibility is that microglia promote angiogenesis in a TREM2 controlled fashion. Indeed pro-angiogenic molecules are highly expressed by GAMs (Brandenburg et al., 2016). Our RNAseq analysis revealed that TREM2 deficiency suppresses the expression of several pro-angiogenic factors, including *Ccl5*, *Ccr5*, *Ccl12*, *Icam1* and *Itgal*. In the similar vein, these angiogenesis related factors are positively correlated with TREM2 expression in CGGA datasets. These TREM2-dependent factors may promote vascularization within tumor environment, which would explain our finding that tumors in WT animals are larger as compared to TREM2 KO. Itgal is a stromal regulator of murine low-grade glioma growth, and it also regulates Ccl5 production (De Andrade Costa et al., 2022). The Ccl5-Ccr5 signaling pathway promotes VEGF-dependent angiogenesis in the osteosarcoma xenograft animal model (Wang et al., 2015). VEGF is an important factor in tumor vascularization, but it has a suppressive effect on the immunologic and pro-angiogenic function of microglia/macrophages in a glioblastoma rodent model (Turkowski et al., 2018). A very interesting candidate is the CXCL2-CXCR2 signaling pathway. In isolated microglia/macrophages from glioma, CXCL2 was strongly up-regulated and showed a higher angiogenic activity than VEGF. Blocking the CXCL2-CXCR2 signaling pathway resulted in considerably diminished glioma sizes (Brandenburg et al., 2016).

In addition to the potential pro-angiogenic role, these factors might also directly stimulate glioma growth. For example, the CCL5-CCR5 axis mediates activation of Akt, and subsequently induces proliferation and invasive behavior of glioma cells (Zhao et al., 2015). Knockdown of Ccl5 promotes glioma cell survival in vitro, and increases the survival in a glioblastoma mouse model (Pan et al., 2017).

We also found a lower density of GAMs in the glioma environment in TREM2 KO animals as compared to WT. Since the number of Ki67^+^ GAMs is similar between these two genotypes, it is likely that TREM2 deficiency impairs the recruitment of monocytes rather than the proliferation of intrinsic microglia. This would be consistent with our observation that the density of Iba1^+^ is similar in WT OBS glioma model compared to TREM2 KO, where the contribution of peripheral monocytes is lacking.

In summary, our study reveals a gender-specific pro-tumorigenic function of TREM2 in glioma. It is due to a complex interplay between microglia which expresses our gene of interest, namely TREM2, and the glioma cells and its vasculature and potentially the invading immune cells. Further studies should address the detailed molecular mechanism of TREM2-dependent angiogenesis in glioma. Thus, TREM2 inhibition has emerged as a novel therapeutic strategy for glioma suppression by slowing vessel growth.

## Acknowledgements

We thank Dr. Marco Colonna at the Washington University School of Medicine provided us the TREM2 KO animal. We thank Dr. Yamei Tan in Sun Yat-Sen Memorial Hospital helped on the TREM2 KO animal sharing.

## Authors’ contributions

XX and HK designed the experiments, analyzed data. XX and HK wrote the manuscript with inputs from XC, as well as inputs from all co-authors. KZ established the *in vivo* model. FZ analyzed immunohistochemistry data. XC performed the *in vivo* model related experiments. YZ and JZ performed *ex vivo* related experiments. XZ analyzed RNA sequencing data. YH and HZ performed experiments using human tissue.

## Data Availability Statement

All data supporting the findings from this study are available from the authors upon reasonable request.

---

**Supplementary Figure S1.**
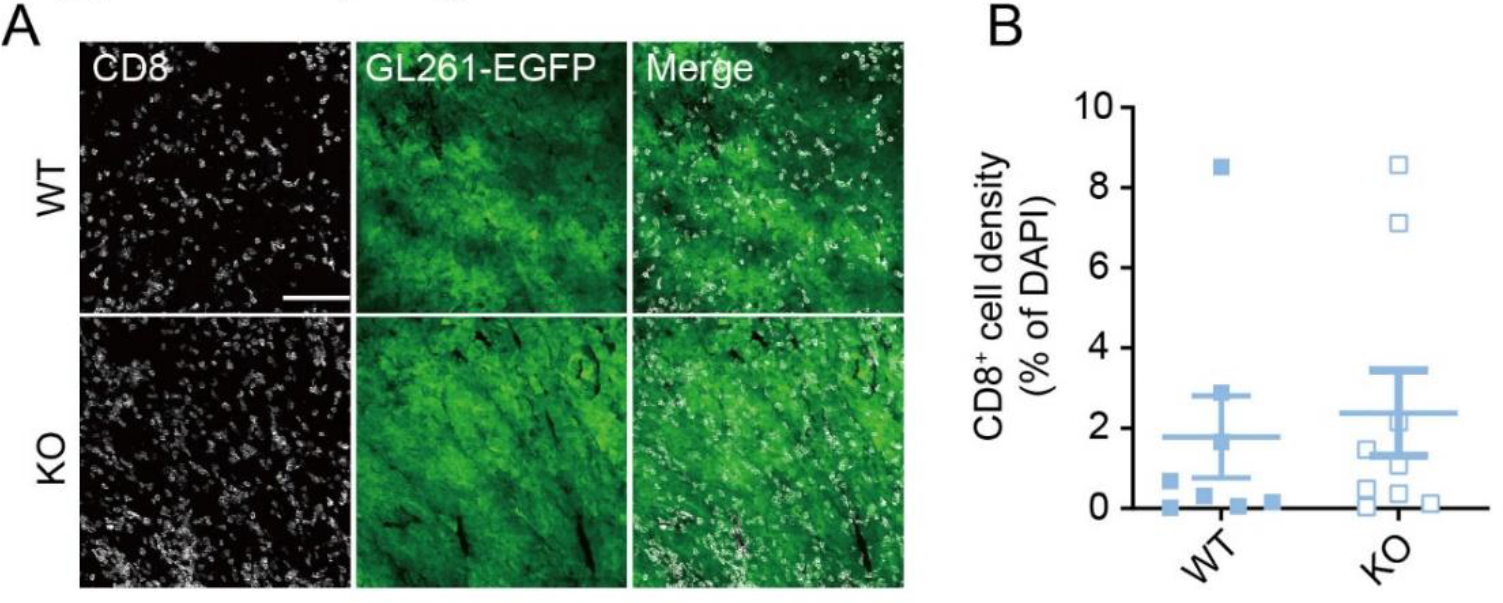
CD8^+^ T cell density is similar between WT and KO **A.** Representative images of CD8 (white), CD31 (red), GL261-EGFP glioma (green) and merged images from WT (top) and TREM2 KO (bottom) animals. (Scale bar, 100 μm) **B.** Quantification of CD8^+^ cells among the DAPI^+^ cells in the glioma tissue of WT and TREM2 KO male mice.

**Supplementary Figure S2.**
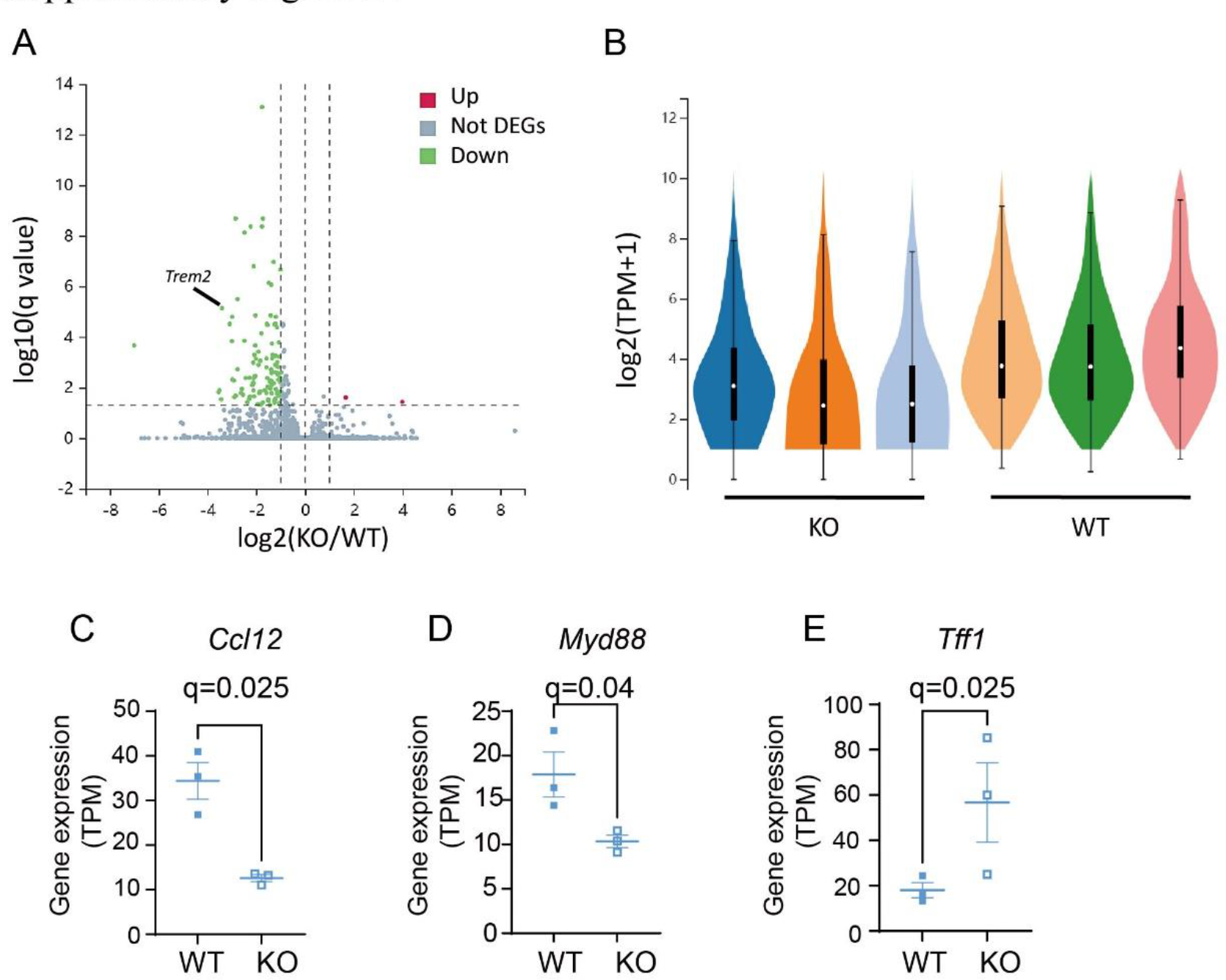
Additional analysis of transcriptomic data for GBM from WT or TREM2 KO mice. A-B. Volcano plot (A, UP: upregulated genes; Down: down regulated genes; Not DEGs: not significantly regulated genes; the value of *Trem2* is indicated) and box plot (B) show the distribution and abundance of 147 DEGs from three different WT and three different KO samples with adjusted p value <0.05 and log2|FC| ≥1. **C-E.** Gene expression (TPM) from WT and TREM2 KO samples of *Ccl12* (C), *Myd88* (D) and *Tff1* (E) (N=3). q values are indicated.

**Supplementary Figure S3.**
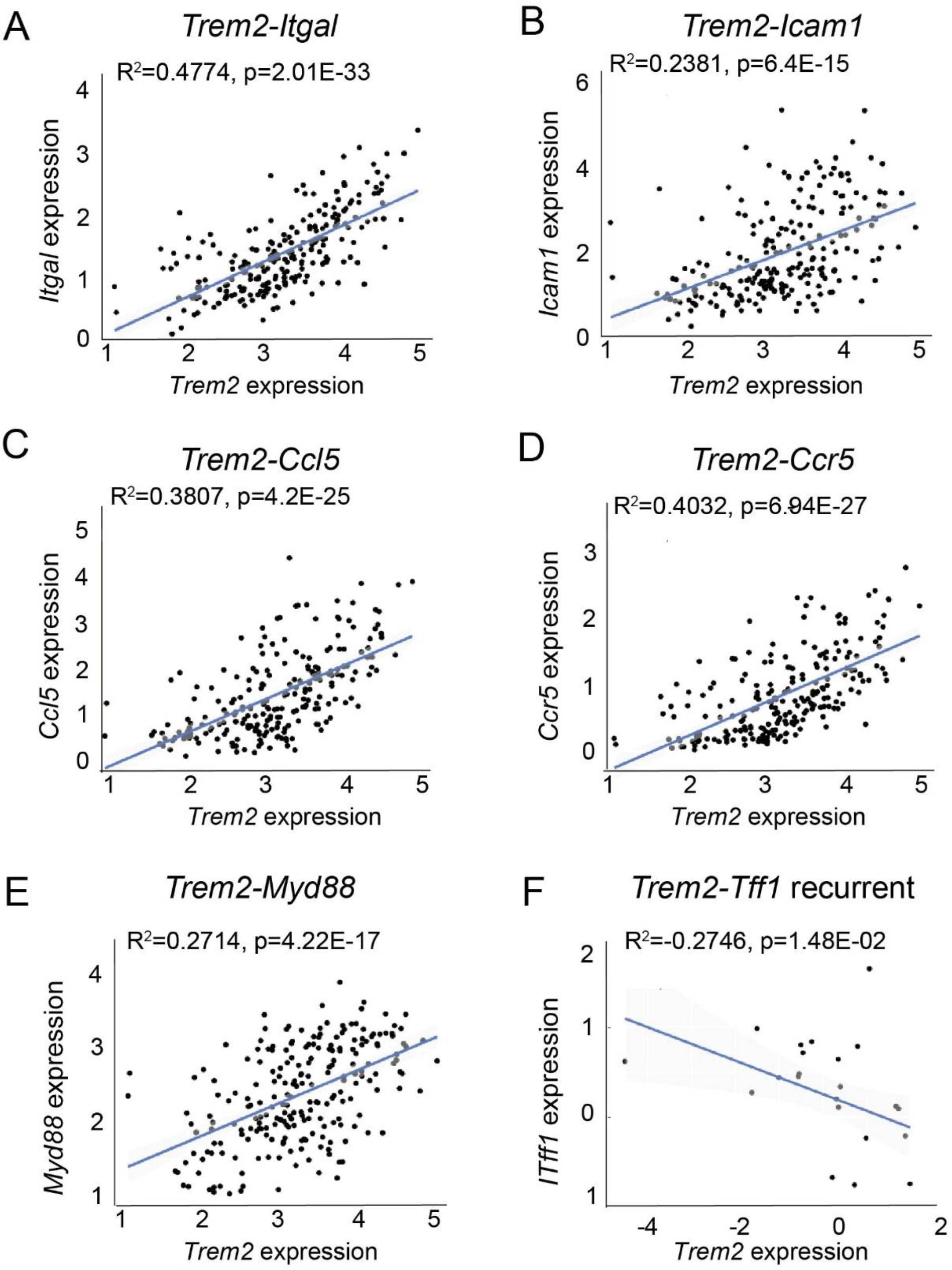
Correlation analysis of *Trem2* with angiogenesis-related genes in glioma patients. Data were collected and analyzed in Chinese Glioma Genome Atlas (CGGA) (http://www.cgga.org.cn/). **A-F.** *Trem2* expression is positively correlated with *Itgal*, *Icam1*, *Ccl5*, *Ccr5*, *Ccl12*, and *Myd88* expression in primary glioma (R= Pearson correlation coefficient; P = P value). **G-H.** *Trem2* expression is negatively correlated with *Tff1* expression in recurrent in microarray dataset.

**Supplementary Figure S4.**
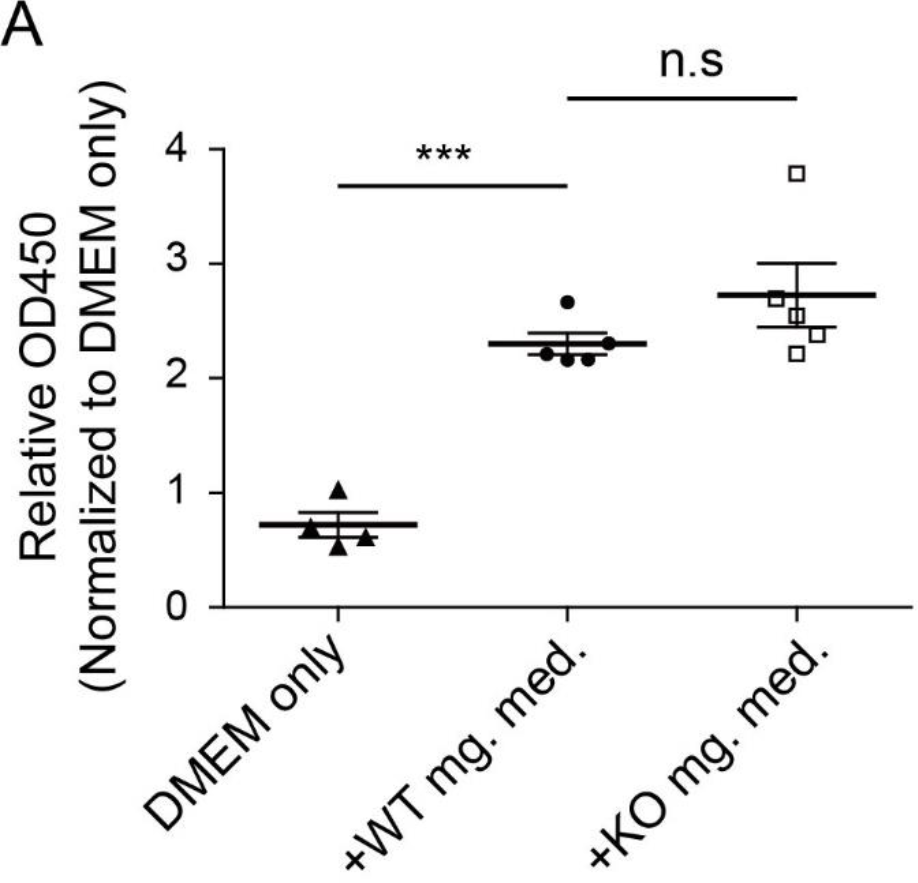
CCK8 cell viability assay **A.** Proliferation rate of GL261 glioma cells was determined using CCK-8 cell counting kit. The absorbance at 450 nm was measured using a microplate reader and normalized to unconditioned DMEM medium. Glioma cells were cultivated in different media conditions as indicated for 48 h. Cultivation in unconditioned DMEM medium was used as a control. WT mg. med: WT primary microglia conditioned DMEM medium; KO mg. med: TREM2 KO primary microglia conditioned DMEM medium. n = 4-6 per group. Data are presented as means ± SEM. One-way ANOVA with Bonferroni corrected multiple comparisons. ***P < 0.001; n.s: non-significant.

## References

De Andrade Costa A, Chatterjee J, Cobb O, Sanapala S, Scheaffer S, Guo X, Dahiya S, Gutmann DH. 2022. RNA sequence analysis reveals ITGAL/CD11A as a stromal regulator of murine low-grade glioma growth. Neuro Oncol 24:14–26.

Binnewies M, Pollack JL, Rudolph J, Dash S, Abushawish M, Lee T, Jahchan NS, Canaday P, Lu E, Norng M, Mankikar S, Liu VM, Du X, Chen A, Mehta R, Palmer R, Juric V, Liang L, Baker KP, Reyno L, Krummel MF, Streuli M, Sriram V. 2021. Targeting TREM2 on tumor-associated macrophages enhances immunotherapy. Cell Rep [Internet] 37:109844. Available from: https://doi.org/10.1016/j.celrep.2021.109844

De Bock K, Cauwenberghs S, Carmeliet P. 2011. Vessel abnormalization: Another hallmark of cancer?. Molecular mechanisms and therapeutic implications. Curr Opin Genet Dev [Internet] 21:73–79. Available from: http://dx.doi.org/10.1016/j.gde.2010.10.008

Bowman RL, Wang Q, Carro A, Verhaak RGW, Squatrito M. 2017. GlioVis data portal for visualization and analysis of brain tumor expression datasets. Neuro Oncol [Internet] 19:139–141. Available from: http://www.ncbi.nlm.nih.gov/pubmed/27630272

Brandenburg S, Müller A, Turkowski K, Radev YT, Rot S, Schmidt C, Bungert AD, Acker G, Schorr A, Hippe A, Miller K, Heppner FL, Homey B, Vajkoczy P. 2016. Resident microglia rather than peripheral macrophages promote vascularization in brain tumors and are source of alternative pro-angiogenic factors. Acta Neuropathol [Internet] 131:365–78. Available from: http://www.ncbi.nlm.nih.gov/pubmed/26718201

Burster T, Gärtner F, Bulach C, Zhanapiya A, Gihring A, Knippschild U. 2021. Regulation of MHC I Molecules in Glioblastoma Cells and the Sensitizing of NK Cells. Pharmaceuticals (Basel) [Internet] 14:1–15. Available from: http://www.ncbi.nlm.nih.gov/pubmed/33800301

Charles NA, Holland EC, Gilbertson R, Glass R, Kettenmann H. 2012. The brain tumor microenvironment. Glia [Internet] 60:502–14. Available from:http://www.ncbi.nlm.nih.gov/pubmed/22379614

Colonna M. 2003. Trems in the immune system and beyond. Nat Rev Immunol 3:445–453.

Davis ME. 2016. Glioblastoma: Overview of disease and treatment. Clin J Oncol Nurs 20:1–8.

Dumas AA, Pomella N, Rosser G, Guglielmi L, Vinel C, Millner TO, Rees J, Aley N, Sheer D, Wei J, Marisetty A, Heimberger AB, Bowman RL, Brandner S, Joyce JA, Marino S. 2020. Microglia promote glioblastoma via mTOR-mediated immunosuppression of the tumour microenvironment. EMBO J [Internet] 39:e103790. Available from: http://www.ncbi.nlm.nih.gov/pubmed/32567735

Fischer I, Gagner JP, Law M, Newcomb EW, Zagzag D. 2005. Angiogenesis in gliomas: Biology and molecular pathophysiology. Brain Pathol 15:297–310.

Guneykaya D, Ivanov A, Hernandez DP, Haage V, Wojtas B, Meyer N, Maricos M, Jordan P, Buonfiglioli A, Gielniewski B, Ochocka N, Cömert C, Friedrich C, Artiles LS, Kaminska B, Mertins P, Beule D, Kettenmann H, Wolf SA. 2018. Transcriptional and Translational Differences of Microglia from Male and Female Brains. Cell Rep [Internet] 24:2773–2783.e6. Available from: http://www.ncbi.nlm.nih.gov/pubmed/30184509

Guneykaya D, Ugursu B, Logiacco F, Popp O, Feiks MA, Meyer N, Wendt S, Semtner M, Cherif F, Gauthier C, Madore C, Yin Z, Çınar Ö, Arslan T, Gerevich Z, Mertins P, Butovsky O, Kettenmann H, Wolf SA. 2023. Sex-specific microglia state in the Neuroligin-4 knock-out mouse model of autism spectrum disorder. Brain Behav Immun 111:61–75.

Gutmann DH, Kettenmann H. 2019. Microglia/Brain Macrophages as Central Drivers of Brain Tumor Pathobiology. Neuron [Internet] 104:442–449. Available from: https://doi.org/10.1016/j.neuron.2019.08.028

Haddad AF, Young JS, Amara D, Berger MS, Raleigh DR, Aghi MK, Butowski NA. 2021. Mouse models of glioblastoma for the evaluation of novel therapeutic strategies. Neuro-oncology Adv [Internet] 3:vdab100. Available from: http://www.ncbi.nlm.nih.gov/pubmed/34466804

Hambardzumyan D, Gutmann DH, Kettenmann H. 2016. The role of microglia and macrophages in glioma maintenance and progression. Nat Neurosci [Internet] 19:20–7. Available from: http://www.ncbi.nlm.nih.gov/pubmed/26713745

Ho VKY, Reijneveld JC, Enting RH, Bienfait HP, Robe P, Baumert BG, Visser O, Dutch Society for Neuro-Oncology (LWNO). 2014. Changing incidence and improved survival of gliomas. Eur J Cancer [Internet] 50:2309–18. Available from: http://dx.doi.org/10.1016/j.ejca.2014.05.019

Hsieh CL, Koike M, Spusta SC, Niemi EC, Yenari M, Nakamura MC, Seaman WE. 2009. A role for TREM2 ligands in the phagocytosis of apoptotic neuronal cells by microglia. J Neurochem [Internet] 109:1144–56. Available from: http://www.ncbi.nlm.nih.gov/pubmed/19302484

Hu F, Dzaye OD, Hahn A, Yu Y, Scavetta RJ, Dittmar G, Kaczmarek AK, Dunning KR, Ricciardelli C, Rinnenthal JL, Heppner FL, Lehnardt S, Synowitz M, Wolf SA, Kettenmann H. 2015. Glioma-derived versican promotes tumor expansion via glioma-associated microglial/macrophages Toll-like receptor 2 signaling. Neuro Oncol [Internet] 17:200–10. Available from: http://www.ncbi.nlm.nih.gov/pubmed/25452390

Huang Y, Zhang Q, Lubas M, Yuan Y, Yalcin F, Efe I, Xia P, Motta E, Buonfiglioli A, Lehnardt S, Dzaye O, Flueh C, Synowitz M, Hu F, Kettenmann H. 2020. Synergistic Toll-like receptor 3/9 signaling affects properties and impairs glioma-promoting activity of microglia. J Neurosci 40:JN-RM-0666-20.

Jackson CM, Choi J, Lim M. 2019. Mechanisms of immunotherapy resistance: lessons from glioblastoma. Nat Immunol [Internet] 20:1100–1109. Available from: http://dx.doi.org/10.1038/s41590-019-0433-y

Katzenelenbogen Y, Sheban F, Yalin A, Yofe I, Svetlichnyy D, Jaitin DA, Bornstein C, Moshe A, Keren-Shaul H, Cohen M, Wang S-Y, Li B, David E, Salame T-M, Weiner A, Amit I. 2020. Coupled scRNA-Seq and Intracellular Protein Activity Reveal an Immunosuppressive Role of TREM2 in Cancer. Cell [Internet] 182:872–885.e19. Available from: https://doi.org/10.1016/j.cell.2020.06.032

Kilian M, Sheinin R, Tan CL, Friedrich M, Krämer C, Kaminitz A, Sanghvi K, Lindner K, Chih Y-C, Cichon F, Richter B, Jung S, Jähne K, Ratliff M, Prins RM, Etminan N, von Deimling A, Wick W, Madi A, Bunse L, Platten M. 2023. MHC class II-restricted antigen presentation is required to prevent dysfunction of cytotoxic T cells by blood-borne myeloids in brain tumors. Cancer Cell [Internet] 41:235–251.e9. Available from: http://www.ncbi.nlm.nih.gov/pubmed/36638785

Kleinberger G, Brendel M, Mracsko E, Wefers B, Groeneweg L, Xiang X, Focke C, Deußing M, Suárez -Calvet M, Mazaheri F, Parhizkar S, Pettkus N, Wurst W, Feederle R, Bartenstein P, Mueggler T, Arzberger T, Knuesel I, Rominger A, Haass C. 2017. The FTD-like syndrome causing TREM2 T66M mutation impairs microglia function, brain perfusion, and glucose metabolism. EMBO J [Internet] 36:1837–1853. Available from: http://emboj.embopress.org/lookup/doi/10.15252/embj.201796516

Kleinberger G, Yamanishi Y, Suárez -Calvet M, Czirr E, Lohmann E, Cuyvers E, Struyfs H, Pettkus N, Wenninger-Weinzierl A, Mazaheri F, Tahirovic S, Lleó A, Alcolea D, Fortea J, Willem M, Lammich S, Molinuevo JL, Sánchez -Valle R, Antonell A, Ramirez A, Heneka MT, Sleegers K, Van Der Zee J, Martin JJ, Engelborghs S, Demirtas-Tatlidede A, Zetterberg H, Van Broeckhoven C, Gurvit H, Wyss-Coray T, Hardy J, Colonna M, Haass C. 2014. TREM2 mutations implicated in neurodegeneration impair cell surface transport and phagocytosis. Sci Transl Med 6.

Klemm F, Maas RR, Bowman RL, Kornete M, Soukup K, Nassiri S, Brouland JP, Iacobuzio-Donahue CA, Brennan C, Tabar V, Gutin PH, Daniel RT, Hegi ME, Joyce JA. 2020. Interrogation of the Microenvironmental Landscape in Brain Tumors Reveals Disease-Specific Alterations of Immune Cells. Cell [Internet] 181:1643–1660.e17. Available from: http://dx.doi.org/10.1016/j.cell.2020.05.007

Lewcock JW, Schlepckow K, Di Paolo G, Tahirovic S, Monroe KM, Haass C. 2020. Emerging Microglia Biology Defines Novel Therapeutic Approaches for Alzheimer’s Disease. Neuron [Internet] 108:801–821. Available from: https://doi.org/10.1016/j.neuron.2020.09.029

Lim E-J, Kang J-H, Kim Y-J, Kim S, Lee S-J. 2022. ICAM-1 promotes cancer progression by regulating SRC activity as an adapter protein in colorectal cancer. Cell Death Dis [Internet] 13:417. Available from: http://www.ncbi.nlm.nih.gov/pubmed/35487888

Lim M, Xia Y, Bettegowda C, Weller M. 2018. Current state of immunotherapy for glioblastoma. Nat Rev Clin Oncol [Internet] 15:422–442. Available from: http://dx.doi.org/10.1038/s41571-018-0003-5

Liu Y-Y, Yao R-Q, Long L-Y, Liu Y-X, Tao B-Y, Liu H-Y, Liu J-L, Li Z, Chen L, Yao Y-M. 2022. Worldwide productivity and research trend of publications concerning glioma-associated macrophage/microglia: A bibliometric study. Front Neurol [Internet] 13:1047162. Available from: http://www.ncbi.nlm.nih.gov/pubmed/36570441

Massey SC, Whitmire P, Doyle TE, Ippolito JE, Mrugala MM, Hu LS, Canoll P, Anderson ARA, Wilson MA, Fitzpatrick SM, McCarthy MM, Rubin JB, Swanson KR. 2021. Sex differences in health and disease: A review of biological sex differences relevant to cancer with a spotlight on glioma. Cancer Lett [Internet] 498:178–187. Available from: https://doi.org/10.1016/j.canlet.2020.07.030

Mazaheri F, Snaidero N, Kleinberger G, Madore C, Daria A, Werner G, Krasemann S, Capell A, Trümbach D, Wurst W, Brunner B, Bultmann S, Tahirovic S, Kerschensteiner M, Misgeld T, Butovsky O, Haass C. 2017. TREM2 deficiency impairs chemotaxis and microglial responses to neuronal injury. EMBO Rep [Internet] 18:1186–1198. Available from: http://embor.embopress.org/lookup/doi/10.15252/embr.201743922

Molgora M, Esaulova E, Vermi W, Hou J, Chen Y, Luo J, Brioschi S, Bugatti M, Omodei AS, Ricci B, Fronick C, Panda SK, Takeuchi Y, Gubin MM, Faccio R, Cella M, Gilfillan S, Unanue ER, Artyomov MN, Schreiber RD, Colonna M. 2020. TREM2 Modulation Remodels the Tumor Myeloid Landscape Enhancing Anti-PD-1 Immunotherapy. Cell [Internet] 182:886–900.e17. Available from: https://doi.org/10.1016/j.cell.2020.07.013

Ostrom QT, Gittleman H, Truitt G, Boscia A, Kruchko C, Barnholtz-Sloan JS. 2018a. CBTRUS Statistical Report: Primary Brain and Other Central Nervous System Tumors Diagnosed in the United States in 2011-2015. Neuro Oncol [Internet] 20:iv1–iv86. Available from: http://www.ncbi.nlm.nih.gov/pubmed/30445539

Ostrom QT, Rubin JB, Lathia JD, Berens ME, Barnholtz-Sloan JS. 2018b. Females have the survival advantage in glioblastoma. Neuro Oncol [Internet] 20:576–577. Available from: http://www.ncbi.nlm.nih.gov/pubmed/29474647

Pan Y, Smithson LJ, Ma Y, Hambardzumyan D, Gutmann DH. 2017. Ccl5 establishes an autocrine high-grade glioma growth regulatory circuit critical for mesenchymal glioblastoma survival. Oncotarget [Internet] 8:32977–32989. Available from: http://www.ncbi.nlm.nih.gov/pubmed/28380429

Roesch S, Rapp C, Dettling S, Herold-Mende C. 2018. When Immune Cells Turn Bad-Tumor-Associated Microglia/Macrophages in Glioma. Int J Mol Sci [Internet] 19. Available from: http://www.ncbi.nlm.nih.gov/pubmed/29389898

Schaff LR, Mellinghoff IK. 2023. Glioblastoma and Other Primary Brain Malignancies in Adults: A Review. JAMA [Internet] 329:574–587. Available from: http://www.ncbi.nlm.nih.gov/pubmed/36809318

Schlepckow K, Monroe KM, Kleinberger G, Cantuti-Castelvetri L, Parhizkar S, Xia D, Willem M, Werner G, Pettkus N, Brunner B, Sülzen A, Nuscher B, Hampel H, Xiang X, Feederle R, Tahirovic S, Park JI, Prorok R, Mahon C, Liang C, Shi J, Kim DJ, Sabelström H, Huang F, Di Paolo G, Simons M, Lewcock JW, Haass C. 2020. Enhancing protective microglial activities with a dual function TREM2 antibody to the stalk region. EMBO Mol Med [Internet] 12:e11227. Available from: https://onlinelibrary.wiley.com/doi/10.15252/emmm.201911227

Shi Y, Huang X, Chen G, Wang Y, Liu Y, Xu W, Tang S, Guleng B, Liu J, Ren J. 2019. miR-632 promotes gastric cancer progression by accelerating angiogenesis in a TFF1-dependent manner. BMC Cancer [Internet] 19:14. Available from: http://www.ncbi.nlm.nih.gov/pubmed/30612555

Silva R, D’Amico G, Hodivala-Dilke KM, Reynolds LE. 2008. Integrins: the keys to unlocking angiogenesis. Arterioscler Thromb Vasc Biol [Internet] 28:1703–13. Available from: http://www.ncbi.nlm.nih.gov/pubmed/18658045

Suffee N, Richard B, Hlawaty H, Oudar O, Charnaux N, Sutton A. 2011. Angiogenic properties of the chemokine RANTES/CCL5. Biochem Soc Trans [Internet] 39:1649–53. Available from: http://www.ncbi.nlm.nih.gov/pubmed/22103502

Turkowski K, Brandenburg S, Mueller A, Kremenetskaia I, Bungert AD, Blank A, Felsenstein M, Vajkoczy P. 2018. VEGF as a modulator of the innate immune response in glioblastoma. Glia [Internet] 66:161–174. Available from: http://www.ncbi.nlm.nih.gov/pubmed/28948650

Turnbull IR, Gilfillan S, Cella M, Aoshi T, Miller M, Piccio L, Hernandez M, Colonna M. 2006. Cutting edge: TREM-2 attenuates macrophage activation. J Immunol [Internet] 177:3520–4. Available from: http://www.ncbi.nlm.nih.gov/pubmed/16951310

VanRyzin JW, Marquardt AE, Pickett LA, McCarthy MM. 2020. Microglia and sexual differentiation of the developing brain: A focus on extrinsic factors. Glia [Internet] 68:1100–1113. Available from: http://www.ncbi.nlm.nih.gov/pubmed/31691400

VanRyzin JW, Pickett LA, McCarthy MM. 2018. Microglia: Driving critical periods and sexual differentiation of the brain. Dev Neurobiol [Internet] 78:580–592. Available from: http://www.ncbi.nlm.nih.gov/pubmed/29243403

Wang S-W, Liu S-C, Sun H-L, Huang T-Y, Chan C-H, Yang C-Y, Yeh H-I, Huang Y-L, Chou W-Y, Lin Y-M, Tang C-H. 2015. CCL5/CCR5 axis induces vascular endothelial growth factor-mediated tumor angiogenesis in human osteosarcoma microenvironment. Carcinogenesis [Internet] 36:104–14. Available from: http://www.ncbi.nlm.nih.gov/pubmed/25330803

Wang S, Mustafa M, Yuede CM, Salazar SV, Kong P, Long H, Ward M, Siddiqui O, Paul R, Gilfillan S, Ibrahim A, Rhinn H, Tassi I, Rosenthal A, Schwabe T, Colonna M. 2020. Anti-human TREM2 induces microglia proliferation and reduces pathology in an Alzheimer’s disease model. J Exp Med [Internet] 217. Available from: http://www.ncbi.nlm.nih.gov/pubmed/32579671

Wu W, Klockow JL, Zhang M, Lafortune F, Chang E, Jin L, Wu Y, Daldrup-Link HE. 2021. Glioblastoma multiforme (GBM): An overview of current therapies and mechanisms of resistance. Pharmacol Res [Internet] 171:105780. Available from: https://doi.org/10.1016/j.phrs.2021.105780

Xiang X, Werner G, Bohrmann B, Liesz A, Mazaheri F, Capell A, Feederle R, Knuesel I, Kleinberger G, Haass C. 2016. TREM2 deficiency reduces the efficacy of immunotherapeutic amyloid clearance. EMBO Mol Med [Internet] 8:992–1004. Available from: http://www.ncbi.nlm.nih.gov/pubmed/26682570

Xie M, Liu YU, Zhao S, Zhang L, Bosco DB, Pang YP, Zhong J, Sheth U, Martens YA, Zhao N, Liu CC, Zhuang Y, Wang L, Dickson DW, Mattson MP, Bu G, Wu LJ. 2022. TREM2 interacts with TDP-43 and mediates microglial neuroprotection against TDP-43-related neurodegeneration. Nat Neurosci 25:26– 38.

Yang W, Warrington NM, Taylor SJ, Whitmire P, Carrasco E, Singleton KW, Wu N, Lathia JD, Berens ME, Kim AH, Barnholtz-Sloan JS, Swanson KR, Luo J, Rubin JB. 2019. Sex differences in GBM revealed by analysis of patient imaging, transcriptome, and survival data. Sci Transl Med [Internet] 11:1–15. Available from: http://www.ncbi.nlm.nih.gov/pubmed/30602536

Yin W, Ping YF, Li F, Lv SQ, Zhang XN, Li XG, Guo Y, Liu Q, Li TR, Yang LQ, Yang K Di, Liu YQ, Luo CH, Luo T, Wang WY, Mao M, Luo M, He ZC, Cao MF, Chen C, Miao JY, Zeng H, Wang C, Zhou L, Yang Y, Yang X, Wang QH, Feng H, Shi Y, Bian XW. 2022. A map of the spatial distribution and tumour-associated macrophage states in glioblastoma and grade 4 IDH-mutant astrocytoma. J Pathol:121–135.

Zhang C, Wang N, Tan H-Y, Guo W, Chen F, Zhong Z, Man K, Tsao SW, Lao L, Feng Y. 2020. Direct inhibition of the TLR4/MyD88 pathway by geniposide suppresses HIF-1α-independent VEGF expression and angiogenesis in hepatocellular carcinoma. Br J Pharmacol [Internet] 177:3240–3257. Available from: http://www.ncbi.nlm.nih.gov/pubmed/32144747

Zhang J, Wang H, Yuan C, Wu J, Xu J, Chen S, Zhang C, He Y. 2022a. ITGAL as a Prognostic Biomarker Correlated With Immune Infiltrates in Gastric Cancer. Front cell Dev Biol [Internet] 10:808212. Available from: http://www.ncbi.nlm.nih.gov/pubmed/35399517

Zhang W, Zhang D, Cheng Y, Liang X, Wang J. 2022b. Runx1 regulates Tff1 expression to expedite viability of retinal microvascular endothelial cells in mice with diabetic retinopathy. Exp Eye Res [Internet] 217:108969. Available from: https://doi.org/10.1016/j.exer.2022.108969

Zhao L, Wang Y, Xue Y, Lv W, Zhang Y, He S. 2015. Critical roles of chemokine receptor CCR5 in regulating glioblastoma proliferation and invasion. Acta Biochim Biophys Sin (Shanghai) [Internet] 47:890–8. Available from: http://www.ncbi.nlm.nih.gov/pubmed/26390883

Zhao Z, Zhang K-N, Wang Q, Li G, Zeng F, Zhang Y, Wu F, Chai R, Wang Z, Zhang C, Zhang W, Bao Z, Jiang T. 2021. Chinese Glioma Genome Atlas (CGGA): A Comprehensive Resource with Functional Genomic Data from Chinese Glioma Patients. Genomics Proteomics Bioinformatics [Internet] 19:1–12. Available from: https://doi.org/10.1016/j.gpb.2020.10.005

